# Transcriptional signatures of progressive neuropathology in transgenic tau and amyloid mouse models

**DOI:** 10.1101/548578

**Authors:** Isabel Castanho, Tracey K. Murray, Eilis Hannon, Aaron Jeffries, Emma Walker, Emma Laing, Hedley Baulf, Joshua Harvey, Andrew Randall, Karen Moore, Paul O’Neill, Katie Lunnon, David A. Collier, Zeshan Ahmed, Michael J. O’Neil, Jonathan Mill

## Abstract

The onset and progression of Alzheimer’s disease (AD) is characterized by increasing intracellular aggregation of hyperphosphorylated tau protein and accumulation of β-amyloid (Aβ) in the neocortex. Despite recent success in identifying genetic risk factors for AD the transcriptional mechanisms involved in disease progression are not fully understood. We used transgenic mice harbouring human tau (rTg4510) and amyloid precursor protein (J20) mutations to investigate transcriptional changes associated with the development of both tau and amyloid pathology. Using highly-parallel RNA sequencing we profiled transcriptional variation in the entorhinal cortex at four time points identifying robust genotype-associated differences in entorhinal cortex gene expression in both models. We quantified neuropathological burden across multiple brain regions in the same individual mice, identifying widespread changes in gene expression paralleling the development of tau pathology in rTg4510 mice. Differentially expressed transcripts included genes associated with familial AD from genetic studies of human patients, and genes annotated to both common and rare variants identified in GWAS and exome-sequencing studies of late-onset sporadic AD. Systems-level analyses identified discrete co-expression networks associated with the progressive accumulation of tau, with these enriched for genes and pathways previously implicated in the neuro-immunological and neurodegenerative processes driving AD pathology. Finally, we report considerable overlap between tau-associated networks and AD-associated co-expression modules identified in the human cortex. Our data provide further support for an immune-response component in the accumulation of tau, and reveal novel molecular pathways associated with the progression of AD neuropathology.

## INTRODUCTION

Alzheimer’s disease (AD) is a chronic neurodegenerative disorder that is characterized by progressive neuropathology and associated cognitive and functional decline^1^. In addition to the loss of synapses and neurons (manifesting as brain atrophy), AD involves two neuropathological hallmarks: the formation of neurofibrillary tangles (NFTs) that result from the intracellular aggregation of hyperphosphorylated tau protein, also a characteristic of other neurodegenerative disorders including frontotemporal dementia (FTD); and the development of senile plaques, which are extracellular deposits mainly composed of β-amyloid (Aβ) protein that have been the focus of extensive efforts in drug discovery. Although these neuropathological signatures of AD have been relatively well characterized in post-mortem human brain tissue, their exact mechanistic role in disease onset and progression remains poorly understood^2^. There have been considerable advances in identifying the genetic risk factors for both familial and sporadic forms of AD; in addition to the autosomal dominant mutations in *APP, PSEN1*, and *PSEN2* that cause early-onset familial AD^3^, the power of genome-wide association studies (GWAS) and exome sequencing in large sample cohorts^4-10^ has been employed to notable success in identifying both common and rare variants associated with late-onset AD. Although the mechanisms by which associated variants mediate disease susceptibility are not well understood, many of the variants are non-coding and hypothesized to involve regulatory disruption to transcriptional networks across affected regions of the brain.

Mouse models of tau and amyloid have played a major role in defining critical pathology-related processes, including facilitating our understanding of the brain’s transcriptional response to the production and gradual deposition of tau and amyloid into tangles and plaques^11^. Recent studies have identified widespread gene expression differences in transgenic mice harbouring a diverse range of AD-associated mutations^12-17^. However, most analyses to date have been undertaken on relatively small numbers of animals and have not attempted to directly relate transcriptional alterations to the progressive burden of pathology in the same mice.

In this study, we systematically assess the transcriptional changes associated with the progression of AD-associated pathology in the mouse brain, using highly-parallel RNA sequencing (RNA-seq) to quantify gene expression changes in the entorhinal cortex, a region characterized by primary and early neuropathology in human AD^18^. We used well-characterized transgenic mouse models of both tau and amyloid pathology, collecting transcriptional data at multiple time-points carefully selected to span from early to late stages of neuropathology in each model (see **Supplementary Figure 1**). First, to investigate transcriptional signatures of progressive tau pathology we used the rTg4510 mouse model, which overexpresses a human mutant (P301L) form of the microtubule-associated protein tau (MAPT)^19,20^. Second, to investigate amyloid pathology we used the J20 mouse line, which expresses a mutant (K670N/M671L and V717F) form of the human amyloid precursor protein (APP)^21,22^. Transcriptional profiles were related to detailed neuropathological measurements of tau and amyloid burden in the same mice, enabling us to directly relate expression changes to the progressive accumulation of neuropathology. We identified robust genotype-associated differences in entorhinal cortex gene expression in both models, and widespread transcriptional changes paralleling the development of tau pathology in rTg4510 mice. Systems-level analyses uncovered discrete co-expression networks associated with the progression of tau pathology that were enriched for genes and pathways implicated in the onset of AD. Finally, we compared these networks to those identified in AD patients, finding considerable overlap with disease-associated co-expression modules identified in the human cortex.

## RESULTS

### Transgenic mice with APP and MAPT mutations are characterized by progressive neuropathology

In both models, right brain hemisphere tissue sections from transgenic (TG) and wild type (WT) littermate control female mice (see **Methods**) were used to immunohistochemically quantify the progression of neuropathology across multiple brain regions (**Supplementary Figure 2a** and **Supplementary Figure 3a**). First, in rTg4510 mice we measured levels of phosphorylated tau (using the antibody PG-5) at 2, 4, 6 and 8 months comparing them to WT controls at the same ages (n = 7-10 animals per group, total n = 74). We identified a dramatic accumulation of tau pathology in the hippocampus (factorial ANOVA, F(3,66) = 69.76, *P* = 1.96E-20) (**Figure 1a-b**). Highly significant increases in phosphorylated tau were also observed within specific sub-regions of the hippocampus and each of the cortical regions we quantified **(Supplementary Figure 2b-i**). Previous studies have shown that rTg4510 mice develop pretangles around 2.5 months of age, with neurofibrillary tangle (NFT) pathology starting in the neocortex and progressing rapidly into the hippocampus and limbic structures with increasing age^19,20,23^. We have also carried out detailed longitudinal analyses of the progression of tau pathology in several parallel cohorts of mice from our rTg4510 colony as previously reported^24,25^. The spread of tau pathology in rTg4510 mice therefore reflects the spread of NFTs with increasing Braak stage in AD^18^. Second, we quantified levels of amyloid pathology (using the antibody b3D6) in J20 mice at ages 6, 8, 10 and 12 months comparing them to WT controls at the same ages (n = 9-10 animals per group, total n = 73). We again identified dramatic increases in pathology in the hippocampus (factorial ANOVA, F(3,68) = 66.85, *P* = 3.00E-20) (**Figure 1c-d**), with a highly significant accumulation of amyloid also observed in each of the cortical regions examined (**Supplementary Figure 3b-d**). These results concur with previous data highlighting progressive deposition of amyloid plaques in the hippocampus and neocortex of J20 mice at 5-7 months, and ubiquitous plaque pathology by 8-10 months of age^22,26^, reflecting the progressive deposition of amyloid seen in individuals with AD^27^. We also quantified neuropathology in the thalamus which, as expected, showed markedly lower levels of both tau (rTg4150, **Supplementary Figure 2j**) and amyloid (J20, **Supplementary Figure 3e**) pathology relative to the other brain regions tested.

**Figure 1.**
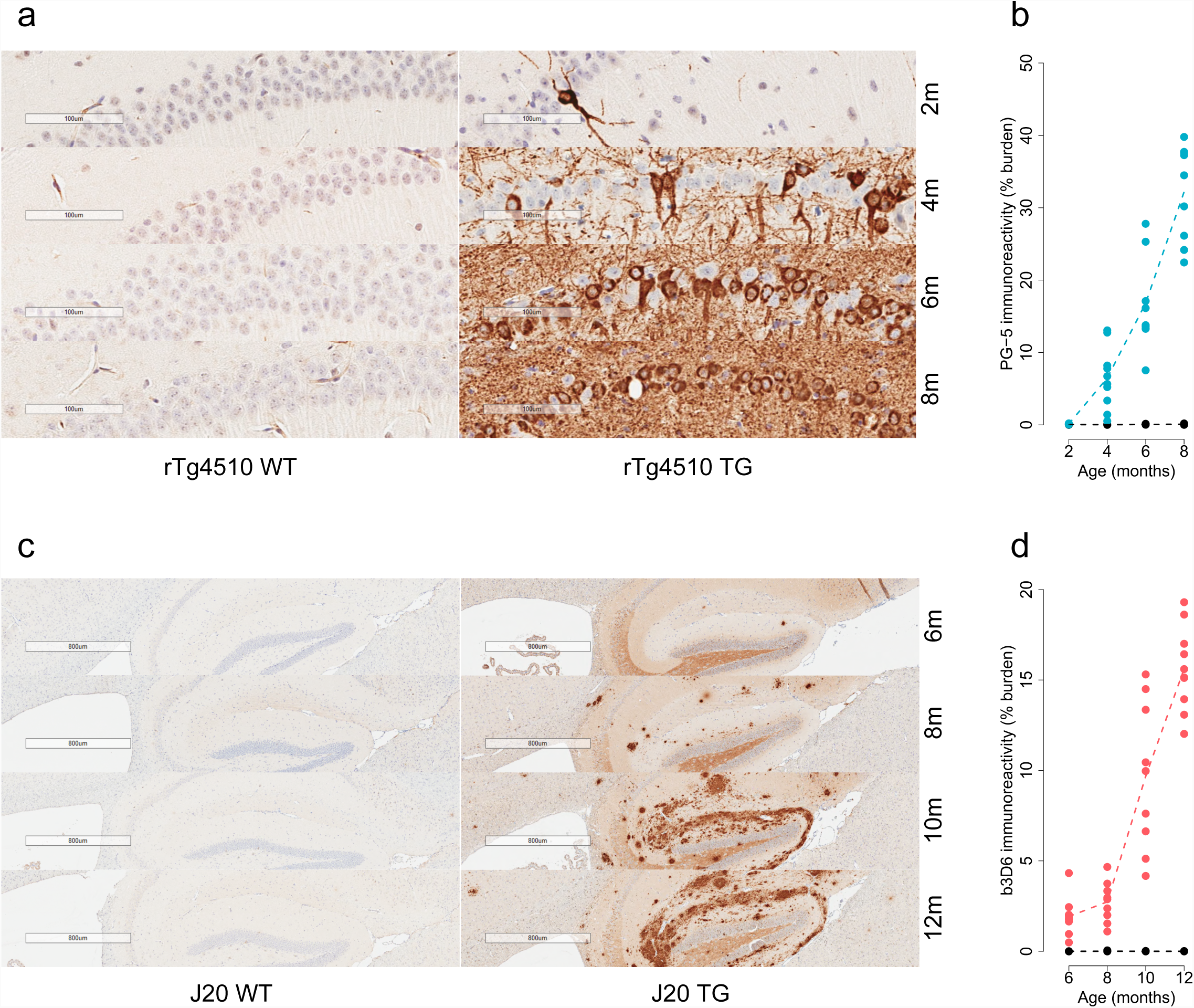
Transgenic models expressing mutant human *MAPT* and *APP* exhibit progressive neuropathology across the hippocampus and cortex. **(a)** Representative immunohistochemistry images from the hippocampus (CA1 subregion) showing the dramatic accumulation of tau pathology (PG-5) in rTg4510 transgenic (TG) mice compared to wild type (WT) control mice at 2, 4, 6 and 8 months of age. **(b)** The quantification of PG-5 immunoreactivity highlighted a striking increase in hippocampal tau in the TG animals (blue) but not WT animals (black) (total n = 73 animals, 8-10 animals per group, factorial ANOVA, F(3,66) = 69.76, *P* = 1.96E-20). **(c)** Representative immunohistochemistry images from the hippocampus showing the dramatic accumulation of amyloid pathology (b3D6) in J20 TG mice compared to WT control mice at 6, 8, 10 and 12 months of age. **(d)** The quantification of b3D6 immunoreactivity highlighted a striking increase in hippocampal amyloid in the TG mice (red) but not WT mice (black) (total n = 77 animals, 9-10 animals per group, factorial ANOVA, F(3,68) = 66.85, *P* = 3.00E-20). Dashed lines represent mean paths of pathological burden across the four age groups.

### The rTg4510 model of tau pathology is characterized by widespread transcriptional differences in the entorhinal cortex

The entorhinal cortex was dissected from the left hemisphere of the brain from each individual mouse (TG and WT) at each of the four time-points. High-quality RNA (mean RIN rTg4510 = 8.9 (SD = 0.2), mean RIN J20 = 8.6 (SD = 0.3)) was isolated from each sample (total n = 121) and used for highly-parallel RNA-seq (see **Methods**). Raw RNA-seq data are available for download from the Gene Expression Omnibus (GEO) database ^28,29^ (accession number GSE125957). After stringent quality control (QC) of the raw RNA-seq data (see **Methods**), we obtained a mean of 18.18 (SD = 3.33) million sequencing reads per sample for the rTg4510 dataset (**Supplementary Table 1**), and a mean of 22.05 (SD = 2.88) million sequencing reads per sample for the J20 dataset (**Supplementary Table 2**), with no difference in read-depth between TG and WT controls (**Supplementary Figure 4**). We quantified read counts for each transcript and evaluated differences in gene expression between TG and WT animals for each model using DESeq2 (see **Methods**). To our knowledge, this represents the most extensive gene expression dataset generated on rodent models of AD pathology, providing excellent power to identify transcriptional variation associated with mutations in *MAPT* and *APP*, and the progressive changes in gene expression accompanying the development of AD pathology in TG mice (**Supplementary Figure 1c**).

Across all samples, striking differences in gene expression were identified in rTg4510 TG animals relative to WT control mice (n = 29 TG, n = 30 WT); gene expression results for all 18,822 detected transcripts are available to download from our online database (www.epigenomicslab.com/ADmice). In total we identified 154 differentially-expressed transcripts at false discovery rate (FDR) < 0.05 (**Figure 2a** and **Supplementary Table 3**). Among these, there was a significant (exact binomial test, n = 154 transcripts, *P* = 0.00014) enrichment of downregulated transcripts (n = 101 (66%) transcripts with reduced expression in TG compared to n = 53 (34%) transcripts with elevated expression in TG). Of note, differences for five of these transcripts are likely to reflect known deletions of the transgene integration sites for the CaMKIIα-tTA (encompassing *Wdr60, Esyt2, Ncapg2*, and *Ptprn2*) and MAPT (encompassing *Fgf14*) transgenes^30^. Given the high homology between transcribed regions of the human and mouse tau gene, we also find highly elevated levels of *Mapt* (Wald statistic = 11.11, log2 fold change = 0.50, FDR = 7.08E-25) (**Supplementary Figure 5a**) confirming stable activation of the MAPT transgene in TG mice; of note, human-specific MAPT sequence domains were only detected in TG RNA-seq datasets (**Supplementary Figure 5b-c**). Furthermore, because the rTg4510 transgene is inserted into the context of two untranslated exons of the mouse prion protein gene (*Prnp*), as expected we observe elevated expression of these *Prnp* domains in TG mice (Wald statistic = 25.40, log2 fold change = 1.54, FDR = 4.88E-138).

**Figure 2.**
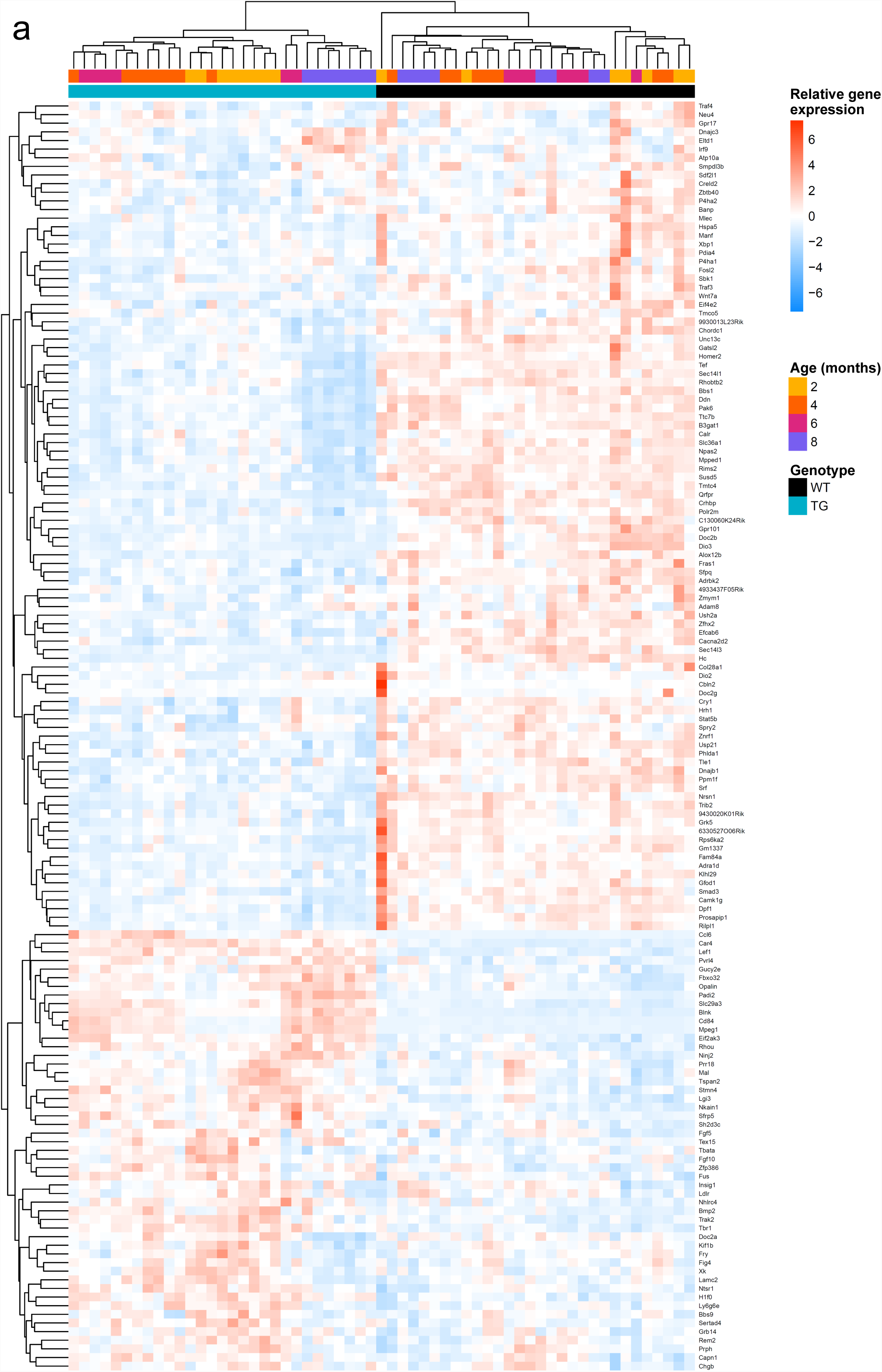
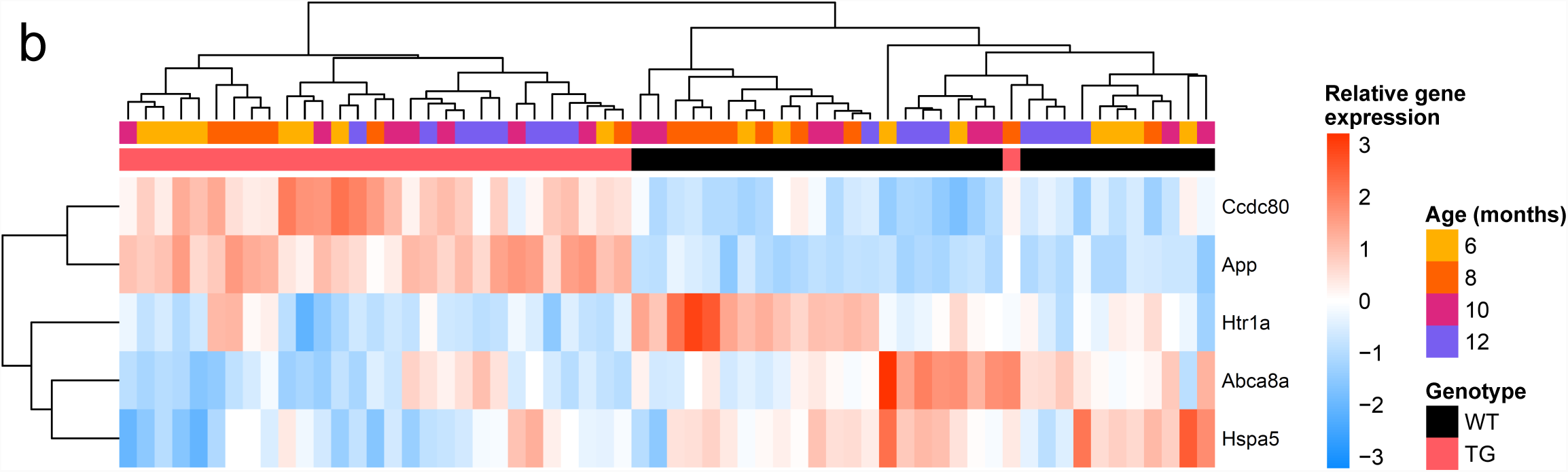
Genotype-associated transcriptional variation robustly discriminates between transgenic and wild type mice. **(a)** Hierarchical clustering of each individual mouse based on expression levels for differentially expressed genes (DEGs) associated with rTg4510 genotype (n = 59 mice (29 TG, 30 WT), 147 transcripts). **(b)** Hierarchical clustering of each individual mouse based on expression levels for differentially expressed genes (DEGs) associated with J20 genotype (n = 62 mice (30 TG, 32 WT), 5 transcripts). Direction of normalized DESeq2 read counts, relative to mean levels of expression across all individual mice (“relative gene expression”), is represented in the heatmaps (scaled) from high (red) to low (blue).

Beyond these expected direct transgene-induced changes, we observed evidence for widespread transcriptional consequences of the rTg4510 genotype. The most significant rTg4510-associated differentially-expressed transcript is *Car4*, which encodes carbonic anhydrase 4 (**Supplementary Figure 6a**, upregulated in TG mice, Wald statistic = 8.36, log2 fold change = 1.11, FDR = 2.41E-13). Other differentially expressed genes in mice carrying the rTg4510 transgene include *Gpr17*, which encodes the G protein-coupled receptor 17 that is involved in regulating oligodendrocyte differentiation and maturation^31^ (**Supplementary Figure 6b**, downregulated in TG mice, Wald statistic = −6.73, log2 fold change = −0.62, FDR = 5.11E-08); *Blnk*, which encodes a cytoplasmic linker protein that plays a critical role in B cell development and is involved in the TREM2 activation pathway^32^ (**Supplementary Figure 6c**, upregulated in TG mice, Wald statistic = 6.48, log2 fold change = 0.80, FDR = 2.12E-07); and *Hspa5* (also known as *Bip* or *Grp78*), which encodes a member of the heat shock protein 70 (HSP70) family that is localized in the lumen of the endoplasmic reticulum (ER) and involved in the folding and assembly of proteins, and has been previously implicated in neuroprotection and AD^33,34^ (**Supplementary Figure 6d**, downregulated in TG mice, Wald statistic = −6.16, log2 fold change = −0.58, FDR = 1.37E-06). Hierarchical clustering of individual mice based on expression levels for genotype-associated transcripts robustly discriminates between rTg4510 and WT groups (**Figure 2a**). Within the rTg4510 TG group, samples also cluster by time-point, suggesting, importantly, that there are progressive changes in gene expression within the mutant mice, and highlighting the value of performing longitudinal analyses.

### The J20 model of amyloid pathology is characterized by differential expression of Ccdc80, Abca8a, Htr1a and Hspa5

Relative to the widespread transcriptional signatures associated with the rTg4510 model, fewer significant expression differences were identified in J20 TG mice compared to WT control mice (n = 30 TG, n = 32 WT); gene expression results for all 18,745 expressed transcripts are available to download from our online database (www.epigenomicslab.com/ADmice). As expected, there was an apparent upregulation of *App* (Wald statistic = 8.55, log2 fold change = 0.66, FDR = 2.37E-13) (**Supplementary Figure 5d**), reflecting the high sequence homology with the human *APP* transgene, and confirming stable activation of the mutant transgene in TG mice; of note, we mapped our RNA-seq reads to human-specific APP sequence domains and only observed signal in TG animals (**Supplementary Figure 5e-f**). In total we identified four additional differentially-expressed transcripts at a stringent false discovery rate (FDR) < 0.05 (**Figure 2b** and **Supplementary Table 4**): *Ccdc80*, encoding a protein involved in cell adhesion and matrix assembly^35^ (upregulated in TG samples, Wald statistic = 6.37, log2 fold change = 0.81, FDR = 1.74E-06); *Abca8a*, encoding a member of the A-subclass of ATP-binding cassette (ABC) transporter family which regulates brain lipid homeostasis and has been implicated in AD^36^ (downregulated in TG samples, Wald statistic = −4.67, log2 fold change = −0.81, FDR = 0.02); *Htr1a*, encoding a major G-protein-coupled serotonin receptor, the 5-HT1A receptor, that is widely expressed in the central nervous system (downregulated in TG samples, Wald statistic = −4.48, log2 fold change = −0.51, FDR = 0.035); and *Hspa5* (downregulated in TG samples, Wald statistic = −4.36, log2 fold change = −0.28, FDR = 0.049) (**Supplementary Figure 7**). Overall, expression of these genotype-associated transcripts discriminates between J20 and WT groups (**Figure 2b**), although, in contrast to the rTg5410 differentially expressed transcripts, there are no clear age effects in the J20 TG mice. Although the transcriptional changes associated with the rTg4510 and J20 genotypes are generally distinct – there is no robust correlation of effect sizes (TG vs WT) between models for differentially expressed transcripts identified in either the rTg4510 (Pearson correlation, r = 0.15, *P* = 0.063, **Supplementary Figure 8a**) or J20 (r = 0.66, *P* = 0.23, **Supplementary Figure 8b**) models – it is noteworthy that *Hspa5* is significantly downregulated (FDR < 0.05) in the same direction in both models (**Supplementary Figure 6d** and **Supplementary Figure 7d**), implicating a role for ER stress in both mouse models.

### Progressive changes in gene expression in the entorhinal cortex mirror the development of neuropathology in animal models of tau and amyloid pathology

Given the progressive accumulation of brain neuropathology in TG mice (**Figure 1**), we next explored temporal changes in gene expression associated with genotype to identify transcriptional signatures paralleling the increases in tau and amyloid pathology in TG mice over time (**Supplementary Figure 1c**). We initially focused on the rTg4510 mice given the clear temporal clustering of samples amongst genotype-associated differentially-expressed transcripts identified in this model (**Figure 2a**). Using an approach designed to identify interactions between genotype (TG vs WT) and age group, we identified 1,762 transcripts (FDR < 0.05) whose expression significantly changed with the progression of tau pathology in rTg4510 mice (**Supplementary Table 5**). Expression differences at these transcripts were found to progressively increase with age relative to baseline (age 2 months) (**Figure 3a**, absolute mean difference (log2 fold change) at 4 months = 0.42; absolute mean difference (log2 fold change) at 6 months = 0.60, absolute mean difference (log2 fold change) at 8 months = 0.78, P < 2.20e-16), paralleling the accumulation of tau pathology in these same individual animals. The top tau-associated differentially-expressed gene in rTg4510 TG mice was *Gfap*, encoding glial fibrillary acidic protein (GFAP), a gene predominantly expressed in both mouse and human astrocytes^37,38^, and known to be upregulated in reactive astrocytes associated with brain pathology^39^. *Gfap* was dramatically upregulated with progressive tau pathology (**Figure 3b**; LRT statistic = 106.321, FDR = 1.28E-18), similar to results from another study reporting age-dependent (12-18 months) upregulation of hippocampal *Gfap* in tau (*CaMKII*-*MAPT* P301L) and amyloid (*APP*/*PSEN1*) mouse models^14^, and paralleling the astrogliosis observed in human AD brain^40,41^. Other top-ranked genes progressively altered in rTg4510 mice were notably enriched for microglial markers previously shown to be upregulated in AD^42,43^, including *Cd68* (**Figure 3c**; LRT statistic = 103.77, FDR = 2.26E-18), *Itgax* (or *Cd11c*) (**Figure 3d**; LRT statistic = 86.85, FDR = 6.54E-15), and *Clec7a* (**Figure 3e**; LRT statistic = 83.20, FDR = 2.97E-14). These genes have all been previously reported to be upregulated in hippocampal tissue from 6-month old rTg4510 female mice^17^, in isolated microglia from rTg4510 mice^25^, in the cortex of amyloid mice at late stages of pathology^15^, and in the neocortex, hippocampus and microglia of mice with amyloid and tau pathology^13,43^. Furthermore, recent transcriptional studies in human brain have shown that microglial gene networks are upregulated in response to AD neuropathology^44^. We used GOseq (see **Methods**) to identify ontological enrichments amongst genes characterized by progressively-altered gene expression in rTg4510 mice, finding highly-significant enrichments for immune-related biological pathways including “immune system process” (FDR = 1.03E-25), “defence response” (FDR = 2.98E-24) and “immune response” (FDR = 4.79E-24) (**Supplementary Table 6**). Given these findings, we next quantified Iba1, a microglia/macrophage-specific calcium-binding protein^45^, in matched tissue sections from the right brain hemisphere (n = 7-10 animals per group, total n = 70), observing a significant increase in all brain regions (hippocampus: factorial ANOVA, F(3,62) = 12.60, *P* = 1.56E-06; cortex: factorial ANOVA, F(3,62) = 18.13, *P* = 1.47E-08; thalamus: factorial ANOVA, F(3,62) = 18.85, *P* = 8.37E-09) (**Supplementary Figure 9**). Together our results reflect the dramatic upregulation of microglial genes observed in studies of other AD rodent models^13-15,43,46,47^, and support a role – either causal or consequential – for dysregulation of the central nervous system (CNS) immune system in the development of AD pathology. Of note, the list of transcripts progressively altered in rTg4510 mice includes genes robustly associated with familial AD from genetic studies of human patients, including *App* (**Supplementary Figure. 10a,** LRT statistic = 13.88, FDR = 0.037) a key driver of amyloid pathology. It also includes genes annotated to both common and rare variants identified in GWAS and exome-sequencing studies of late-onset sporadic AD (LOAD), including *Trem2* (LRT statistic = 43.82, FDR = 3.73E-07), *Pld3* (LRT statistic = 36.80, FDR = 5.80E-06), *Frmd4a* (LRT statistic = 27.81, FDR = 0.00022), *Clu* (LRT statistic = 27.73, FDR = 0.00023), *Apoe* (LRT statistic = 22.99, FDR = 0.0014), *Picalm* (LRT statistic = 21.37, FDR = 0.0025), *Cd33* (LRT statistic = 27.32, FDR = 0.00026), and *Abi3* (LRT statistic = 17.10, FDR = 0.012) (**Supplementary Figure 10b-i**).

**Figure 3.**
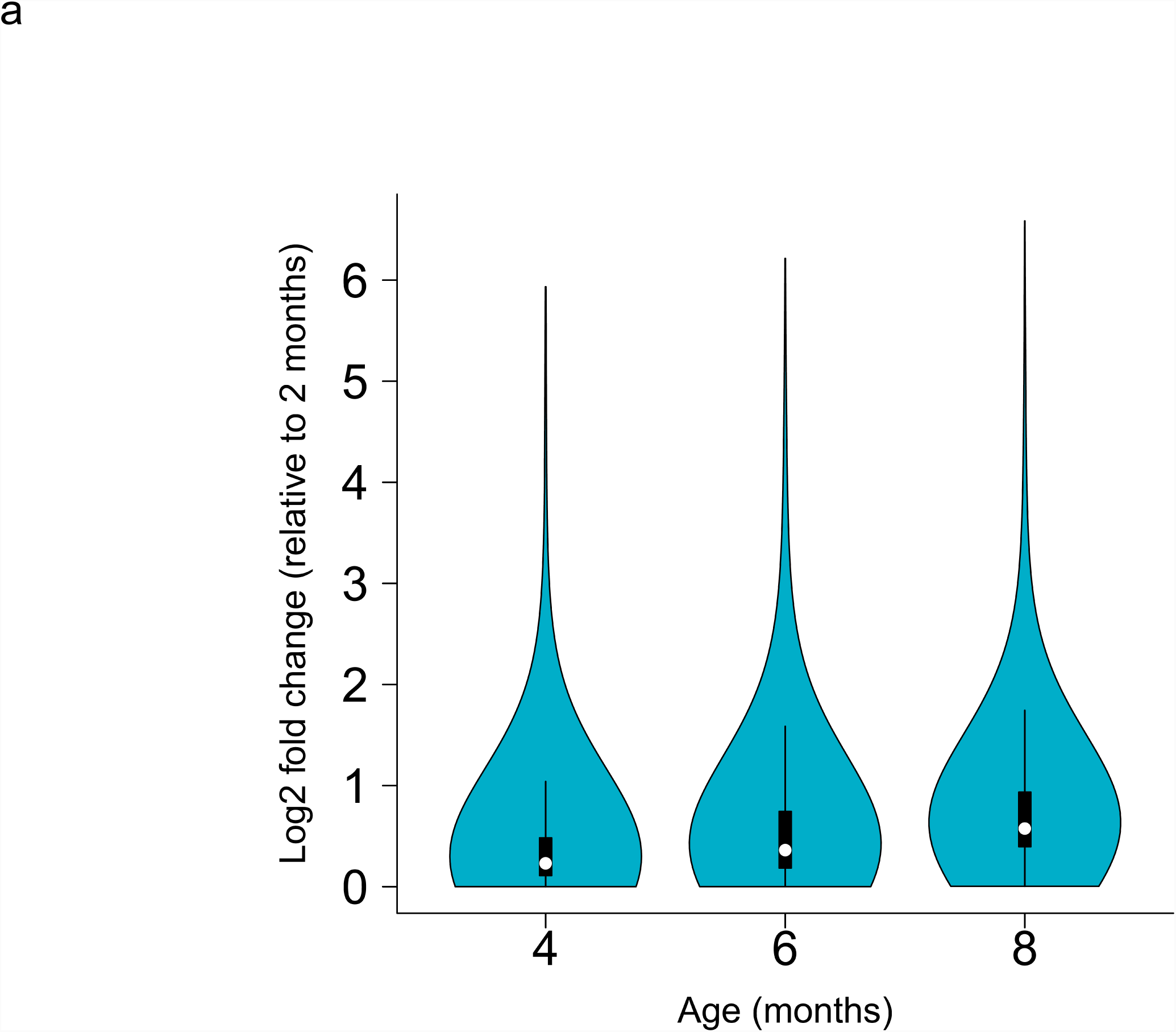
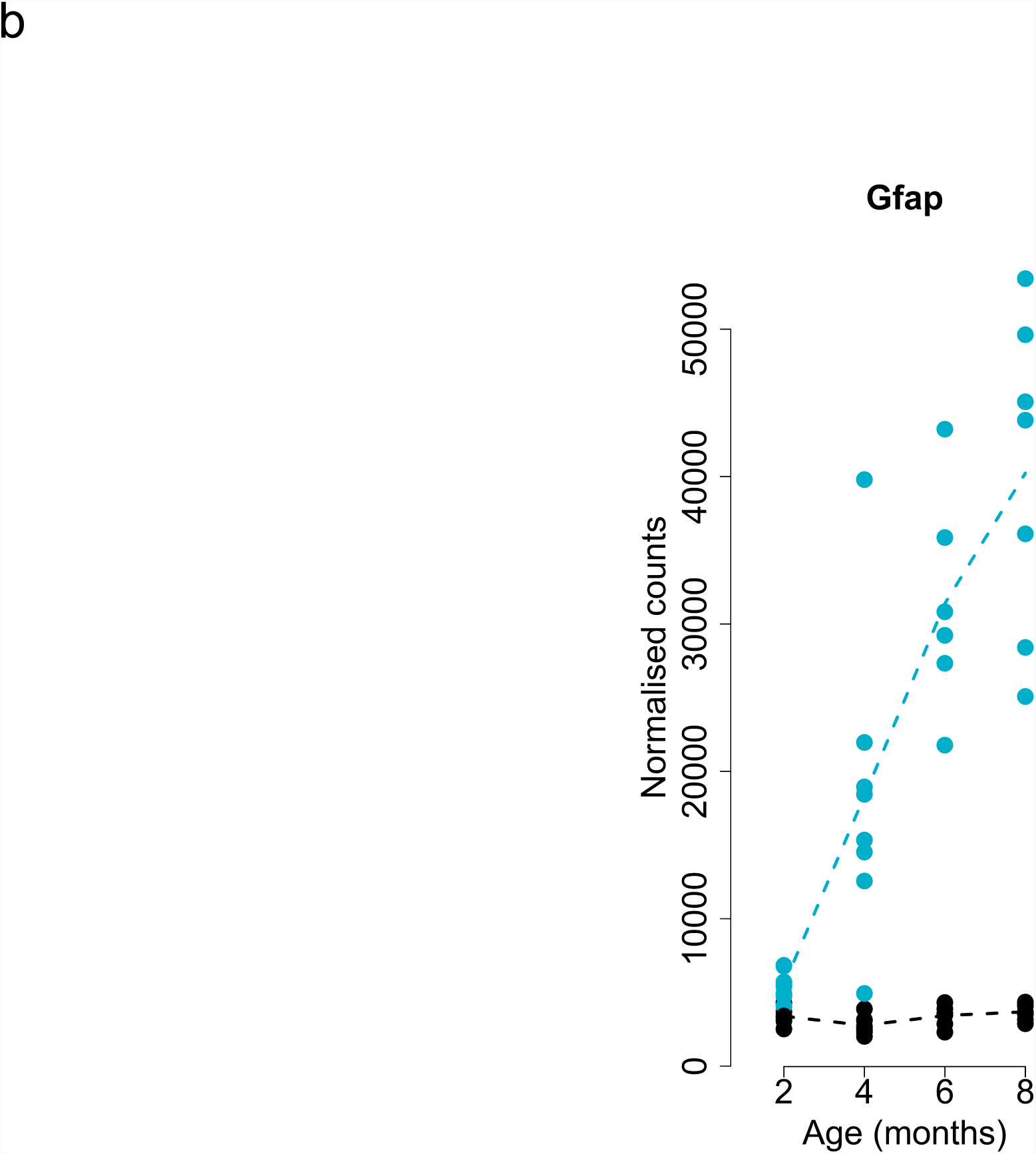
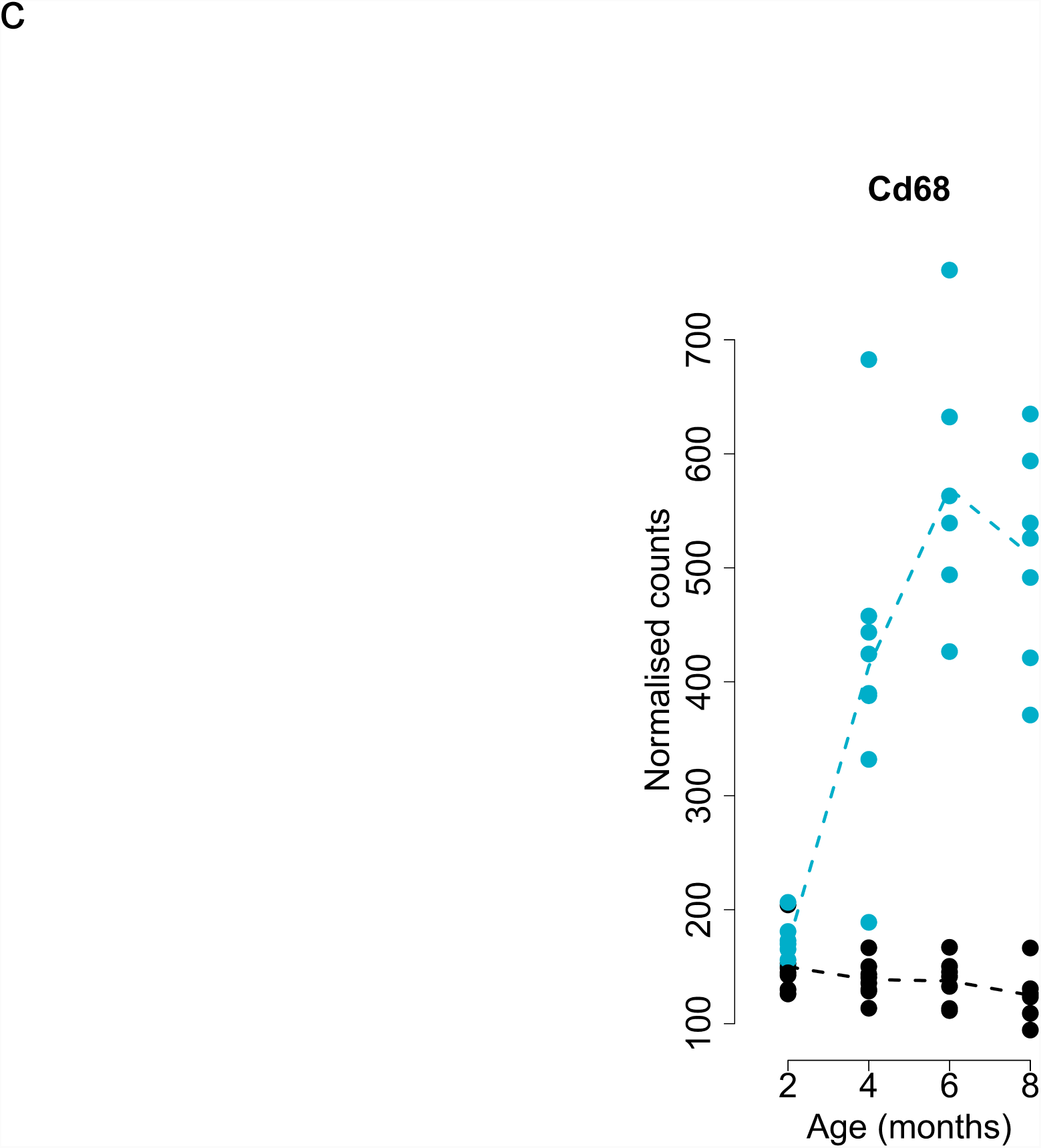
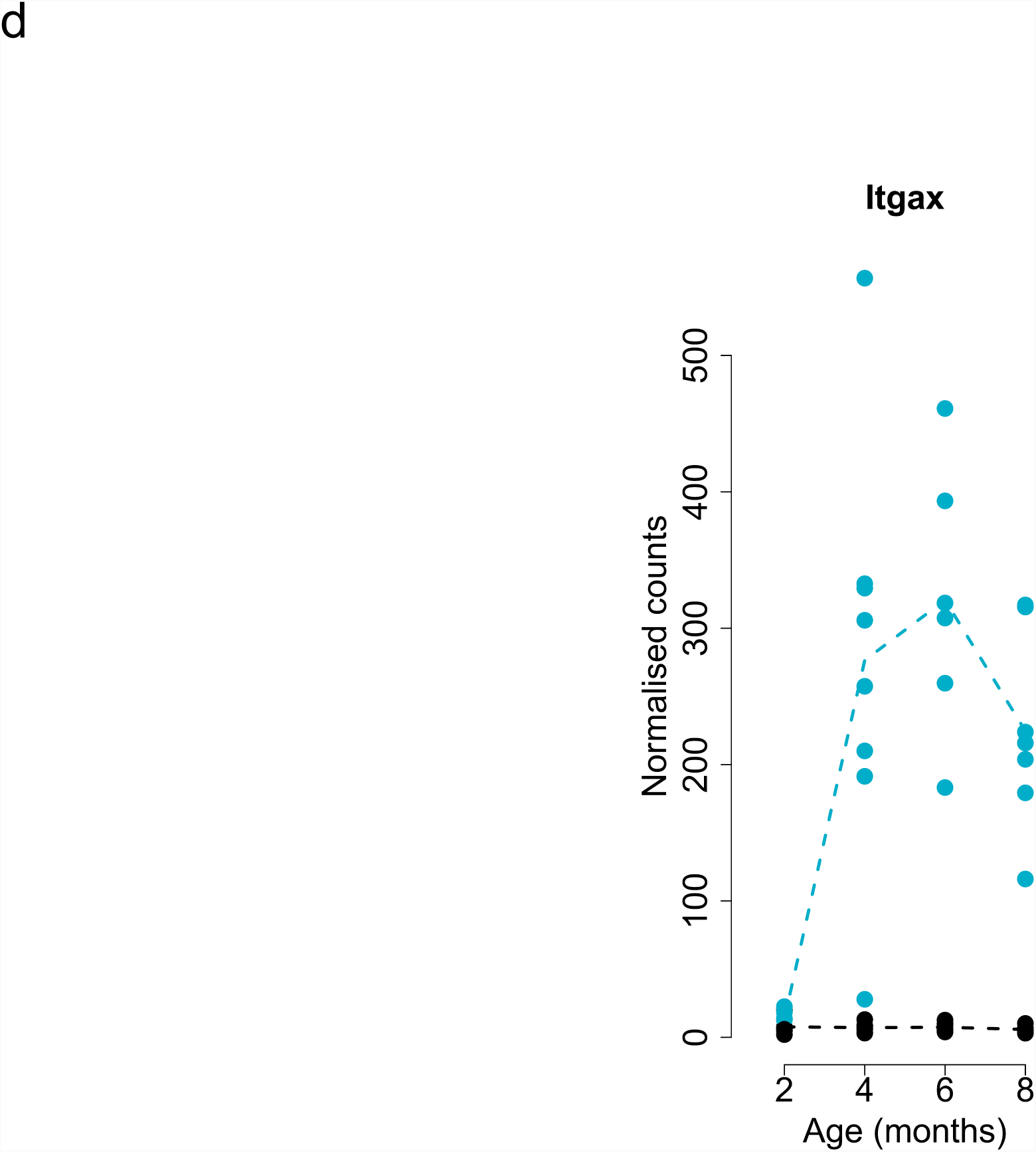
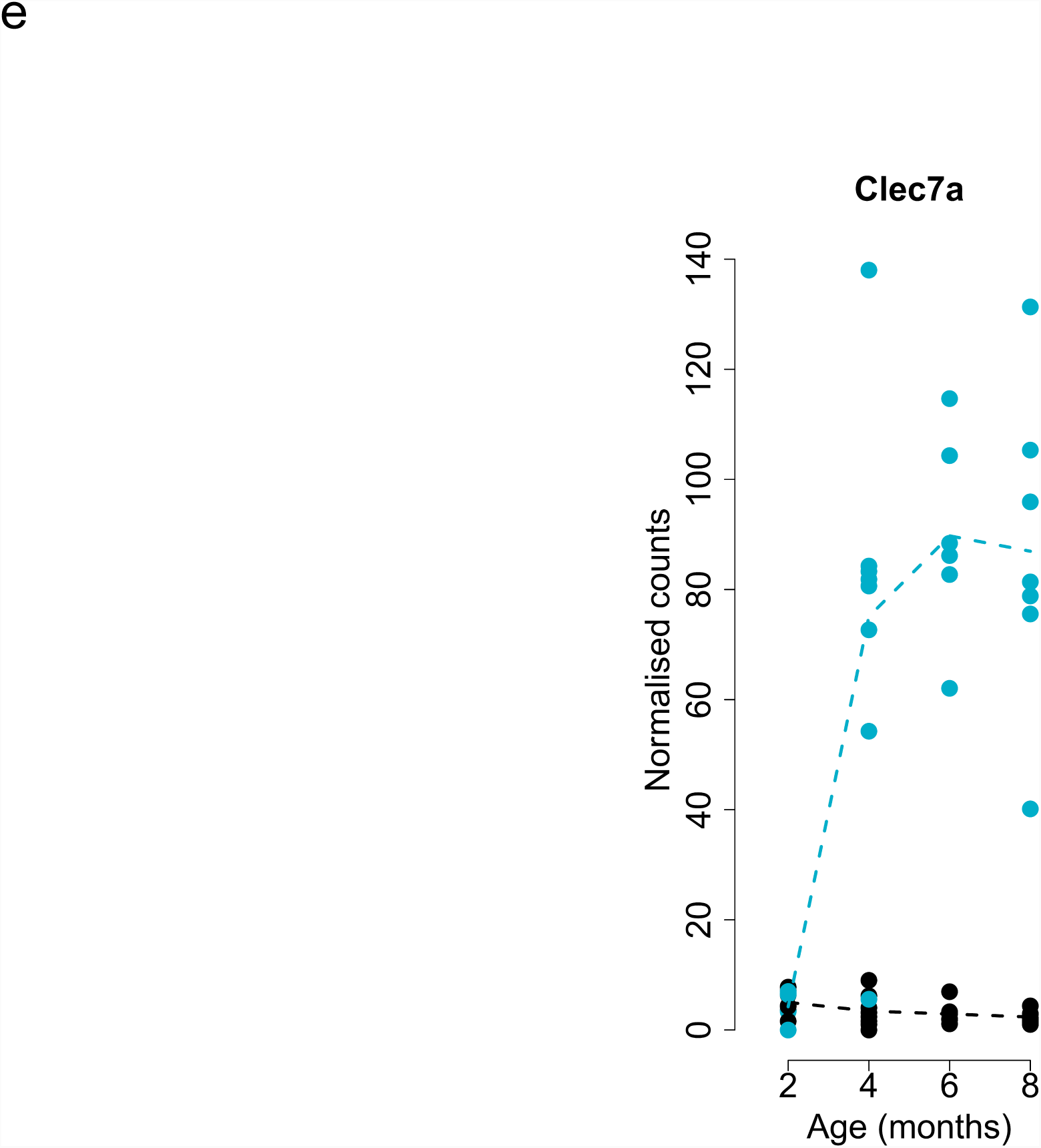
Transcriptional trajectories associated with the accumulation of tau pathology in rTg4510 mice. **(a)** Violin plot showing increasing absolute effect size (Log2 fold change) across age groups for transcripts characterized by significant (FDR < 0.05, n = 1,762 transcripts) temporal changes in gene expression associated with genotype (Mann-Whitney U test, W (4-6 months) = 1182700, P (4-6 months) < 2.20e-16, W (6-8 months) = 1031000, P (6-8 months) < 2.20e-16, W (4-8 months) = 679520, P (4-8 months) < 2.20e-16). White dots represent the median absolute fold-change. Shown are individual plots for: **(b)** *Gfap* (Likelihood-ratio test, LRT statistic = 106.32, log2 fold change (2-8 months) = 2.75, FDR = 1.28E-18), **(c)** *Cd68* (Likelihood-ratio test, LRT statistic = 103.77, log2 fold change (2-8 months) = 1.85, FDR = 2.26E-18), **(d)** *Itgax* (Likelihood-ratio test, LRT statistic = 86.85, log2 fold change (2-8 months) = 4.42, FDR = 6.54E-15), and **(e)** *Clec7a* (Likelihood-ratio test, LRT statistic = 83.20, log2 fold change (2-8 months) = 5.37, FDR = 2.97E-14). Normalized RNA-seq read counts were obtained using *DESeq2*. Dashed lines represent mean paths for each time point. rTg4510 transgenic (TG, blue) female mice compared to wild type (WT, black) littermate control mice. Total n = 59 animals (6-8 animals per group).

In contrast to the dramatic and progressive changes in gene expression identified in rTg4510 mice, fewer significant temporal transcriptional differences associated with the progression of amyloid pathology were identified in the J20 mice; in total we identified five transcripts (*Cst7, Wdfy1, Grxcr2, Itgax*, and *Ifitm1*) whose expression profile significantly changed (FDR < 0.05) with the progression of amyloid pathology (**Supplementary Table 7**). The relatively low number of significantly-altered genes in J20 mice potentially reflects the slower and later accumulation of pathology in these mice compared to rTg4510 mice. Previous work has shown relatively limited amyloid pathology in J20 entorhinal cortex even at 14 months of age^48^, with neuronal cell loss varying by brain region^49^. Nevertheless, we found that effect sizes for the 1,762 transcripts identified as being progressively dysregulated in rTg4510 mice were significantly correlated across both models, suggesting some common molecular signals associated with both tau and amyloid pathology (**Figure 4a**; Pearson correlation, r = 0.46, *P* = 1.50E-92; exact binomial test, n = 1762 transcripts, *P* = 1.97e-05). Interestingly, two genes identified as being associated with progressive tau pathology in rTg4510 mice were also significantly associated with amyloid pathology in J20 mice (*Cst7* and *Itgax* (**Figure 4b-d** and **Supplementary Figure 11**)). Like *Itgax, Cst7* has been shown to be a marker for activated microglia and upregulated in AD pathology^43,50^; of note, *Cst7* was previously reported to be the top upregulated gene in cortex samples from 12 month-old APP NL-G-F knock-in mice^12^.

**Figure 4.**
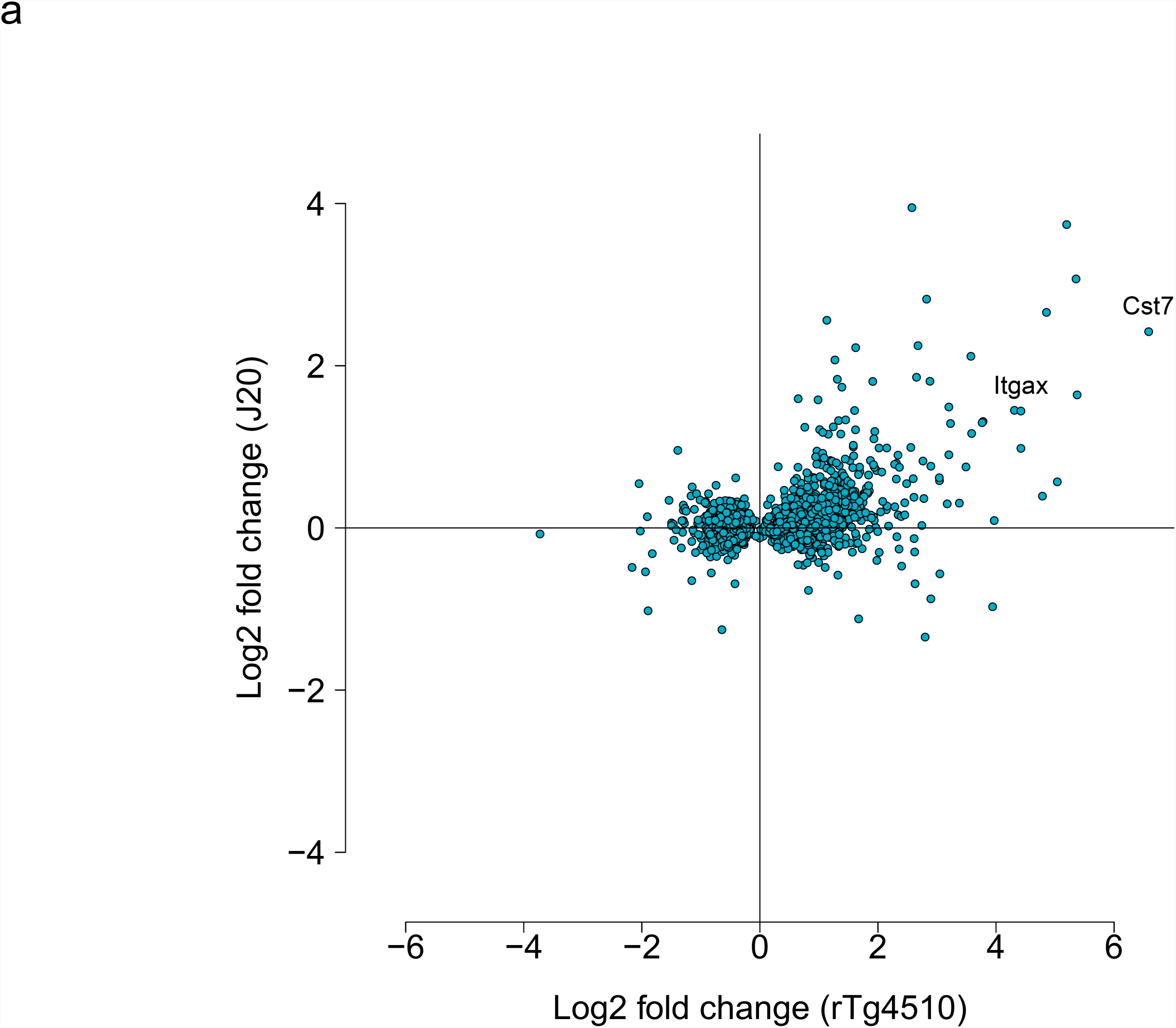
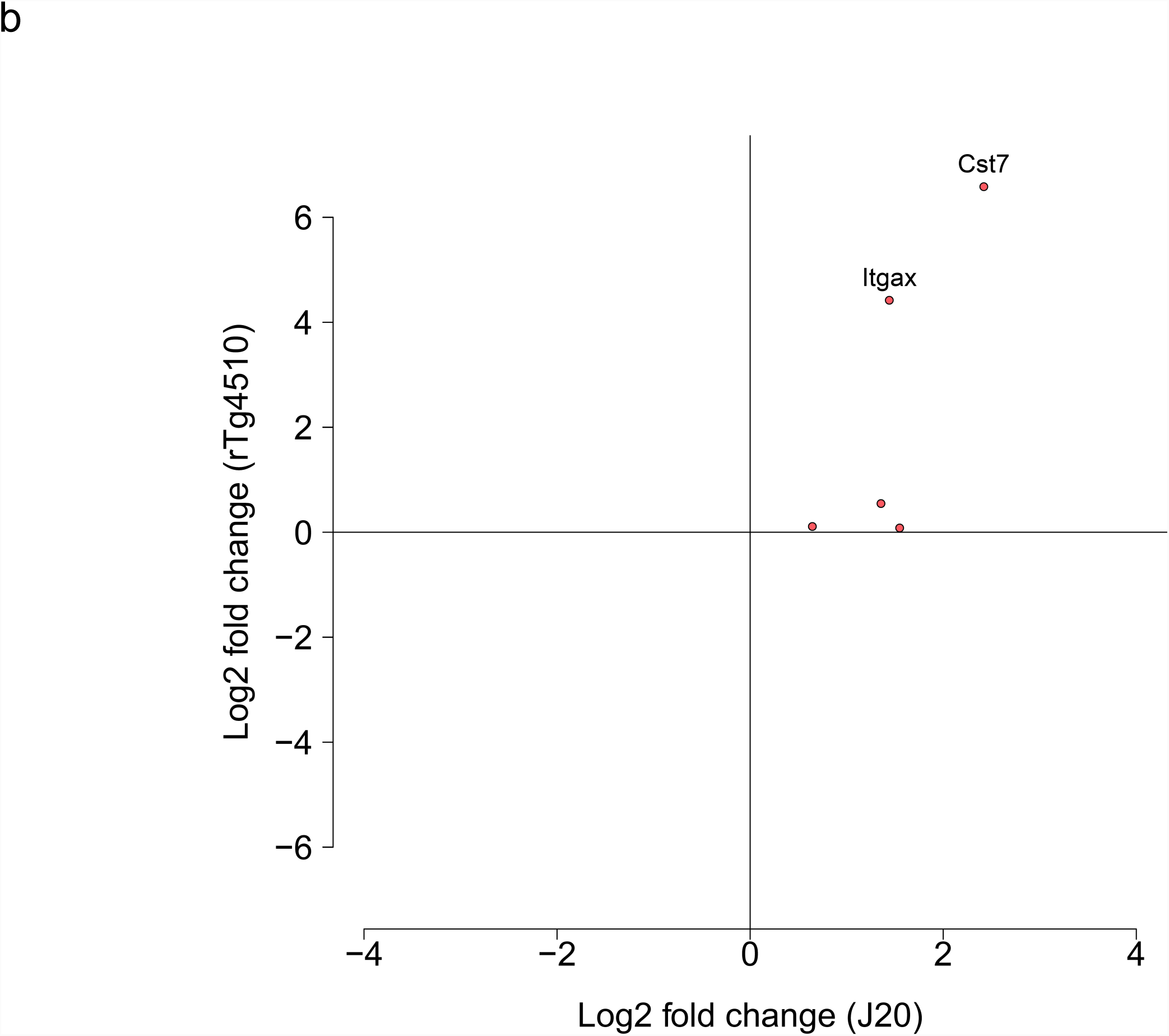
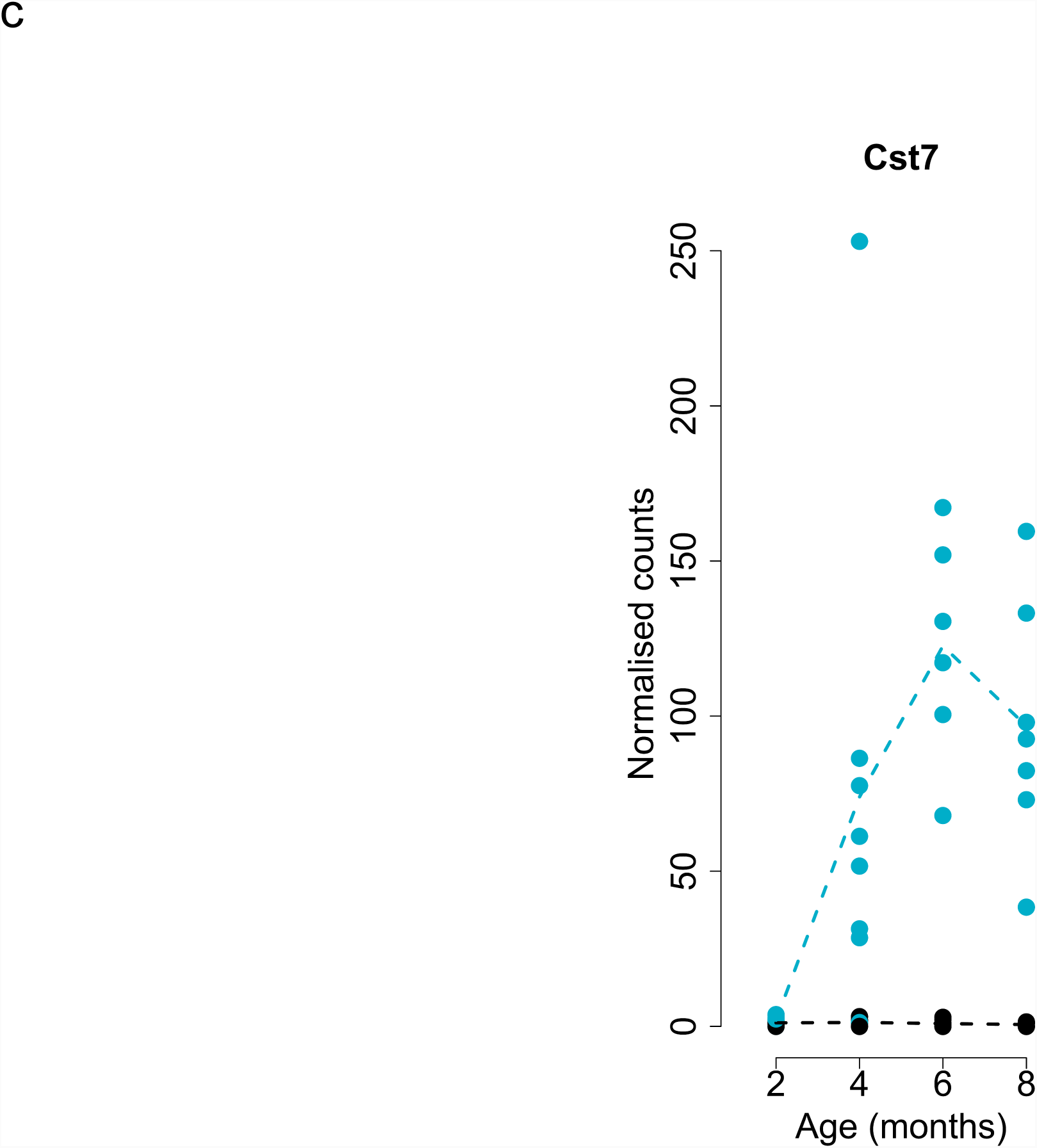
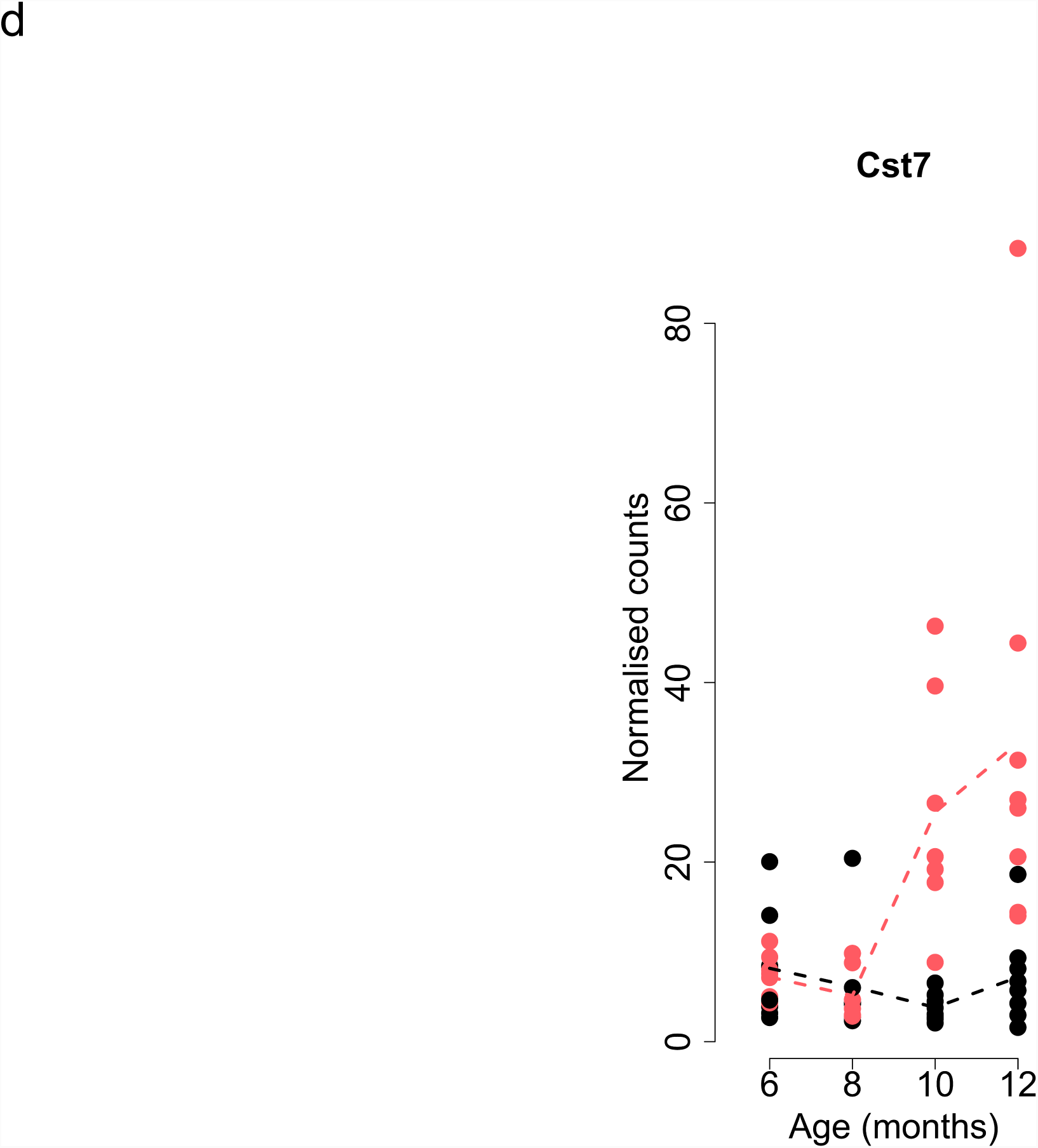
Effect sizes at differentially expressed transcripts associated with the progression of tau in rTg4510 mice are correlated with those in another transgenic model of tau. **(a)** Positive correlation for effect size (Log2 fold change from latest time point compared to baseline) for significant transcripts in rTg4510 mice (Pearson correlation, r = 0.46, *P* = 1.50E-92; exact binomial test, n = 1762 transcripts, P = 1.97e-05). **(b)** Two transcripts (*Cst7* and *Itgax*) were significantly associated with the progression of both tau (Tg4510) and amyloid (J20) pathology (Pearson correlation, r = 0.77, *P* = 0.13; exact binomial test, n = 5 transcripts, P = 0.13). **(c)** *Cst7* gene expression in rTg4510 mice (Total n = 59 animals, 6-8 animals per group, Likelihood-ratio test, LRT statistic = 36.10, log2 fold change (2-8 months) = 6.59, FDR = 7.71E-06). rTg4510 transgenic (TG, blue) female mice compared to wild type (WT, black) littermate control mice. **(d)** *Cst7* gene expression in the J20 mice (Total n = 62 animals, 6-8 animals per group, Likelihood-ratio test, LRT statistic = 37.37, log2 fold change (6-12 months) = 2.42, FDR = 0.00072). J20 transgenic (TG, red) female mice compared to wild type (WT, black) littermate control mice. *Itgax* gene expression in rTg4510 and J20 mice is show in **Figure 3d** and **Supplementary Figure 11**, respectively. Normalized counts were obtained using *DESeq2*. Dashed lines represent mean paths for each time point.

### Transcriptional changes identified in rTg4510 mice reflect those observed in other models of tau pathology

A number of recent studies have described further evidence for differential gene expression in transgenic models of familial AD gene mutations^12-15,17,25,43^. We therefore explored hippocampal RNA-seq data from two other transgenic models (TAU (CaMKII-MAPTP301L) and TAS10 (SwAPP, K670N/M671L)) downloaded from the Mouseac database^14,51^ (www.mouseac.org) to identify consistencies in the transcriptional signatures between different models of tau and amyloid pathology. Effect sizes for transcripts identified as associated with rTg4510 genotype and also present in the Mouseac TAU RNA-seq dataset (n = 138) were significantly correlated between the two models (r = 0.33, *P* = 7.7E-05). Despite this consistency in effect sizes, many of the differentially expressed genes associated with rTg4510 genotype were not statistically replicated in the TAU model (**Supplementary Figure 12** and **Supplementary Table 8**), although this likely reflects the distinct genetic background of the different transgenic lines and the modest power to detect effects given the small number of samples profiled in the Mouseac dataset (n = 49 RNA-seq samples, 1-4 animals per age group, after filtering for samples with complete phenotypic data). Differential expression of *Gpr17* was associated with TAU genotype (Bonferroni-corrected P < 0.00035), although the exact profile for this gene differed to that observed in the rTg4510 mice (**Supplementary Figure 13**). This putative receptor for leukotrienes, uracil nucleotides and/or oxysterols may warrant further investigation as its expression is associated with damage to neural tissue including white matter^52^. As expected, given the limited evidence for consistency in genotype effects between rTg4510 and J20 mice, there was no correlation between effects observed in rTg4510 and TAS10 mice for the 145 rTg4510-associated genes present in both datasets. In contrast, association statistics for the 1640 transcripts identified as being progressively altered with age in rTg4510 mice and also present in the Mouseac datasets (**Supplementary Table 9** and **Supplementary Table 10**) were significantly correlated with those for the same genes in both TAU (r = 0.46, *P* = 1.2e-86) and TAS10 (r = 0.23, *P* = 3.9E-21) transgenic mice (**Supplementary Figure 14**). Given the small number of progressive alterations observed in J20 mice, it was not possible to systematically explore overlaps between differentially regulated genes in this model and the two Mouseac models. Of note, however, the two genes identified as being temporally-altered in both rTg4510 and J20 mice – *Cst7* and *Itgax* – were both similarly altered in both the TAU and TAS10 models (**Supplementary Figure 15**).

### Gene co-expression networks associated with the progression of tau pathology are enriched for functional pathways related to AD including synaptic transmission, the immune system, and glial cell activation

Given the dramatic transcriptional changes identified in the rTg4510 mice, we next used weighted gene correlation network analysis (WGCNA) (see **Methods**) to identify discrete co-expression modules and describe systems-level transcriptional variation associated with rTg4510 genotype and the progression of tau pathology. We constructed co-expression networks using entorhinal cortex RNA-seq data from rTg4510 TG and WT mice (n = 58 mice), identifying 18 discrete co-expression modules (**Supplementary Figure 16**). Next, we used a linear regression model (see **Methods**) and identified six co-expression modules (here named as “salmon”, “turquoise”, “purple”, “yellow”, “light-cyan”, and “red”) that were significantly (Bonferroni corrected, *P* < 0.0028) associated with rTg4510 genotype (**Supplementary Figure 16, Supplementary Figure 17** and **Supplementary Table 11**). Strikingly, these tau-associated co-expression modules are highly enriched for molecular functions and biological pathways directly related to AD. The red module, for example, which was down-regulated in TG mice compared to WT mice (β = −0.18, *P* = 1.43E-10), is highly enriched for functional pathways involved in synaptic transmission (**Supplementary Table 12**). The turquoise module, which was up-regulated in TG mice compared to WT mice (β = 0.18, *P* = 3.04E-10), is enriched for pathways involved in activation of the immune system (**Supplementary Table 13**). The salmon module, which was consistently up-regulated in TG mice compared to WT mice (β = 0.14, *P* = 3.58E-06), is enriched for genes involved in myelination and glial cell activation (**Supplementary Table 14**). The purple module, which was down-regulated in TG mice compared to WT mice (β = −0.13, *P* = 0.00012), is enriched for pathways related to cellular component disassembly (**Supplementary Table 15**). Finally, the yellow module, which was down-regulated in TG mice compared to WT mice (β = −0.10, *P* = 0.0015), is enriched for pathways related to mitochondria and synaptic processes (**Supplementary Table 16**). The module eigengenes for three of these co-expression modules (turquoise, yellow and red) were characterized by a significant interaction between genotype and age in rTg4510 mice (**Supplementary Figure 16** and **Supplementary Table 11**), suggesting that they are temporally linked to the development of tau pathology in TG mice (**Figure 5a-5c**). The turquoise module becomes increasingly up-regulated with the development of tau pathology (β = 0.28, *P* = 4.23E-06), the red module becomes increasingly down-regulated with the development of tau pathology (β = −0.21, *P* = 0.0022), and the yellow module becomes down-regulated specifically during the later stages of tau pathology (β = −0.29, *P* = 0.0018). Using the matched immunohistochemistry data generated across multiple brain regions for each mouse we were able to explore the relationship between co-expression modules and actual tau pathology in rTg4510 mice, confirming that the turquoise, yellow, and red modules are robustly associated with the accumulation of tau across the brain (**Supplementary Figure 18**). The association with pathology was particularly strong in highly affected brain regions such as the hippocampus (**Figure 5d-5f**, **Supplementary Figure 19**, and **Supplementary Figure 20a-c**), in which the module eigengene for the turquoise module is positively correlated with levels of tau in TG mice (r = 0.85, *P* = 1.20E-16), and those for the yellow and red modules are negatively correlated with levels of tau in TG mice (yellow: r = - 0.63, *P* = 2.00E-07, red: r = −0.79, *P* = 4.61E-13). Although these co-expression modules were also correlated with measures of tau pathology in the thalamus, the magnitude of effects was much lower, reflecting the later and less aggressive accumulation of tau in this region of the brain (**Supplementary Figure 20d-f**).

**Figure 5.**
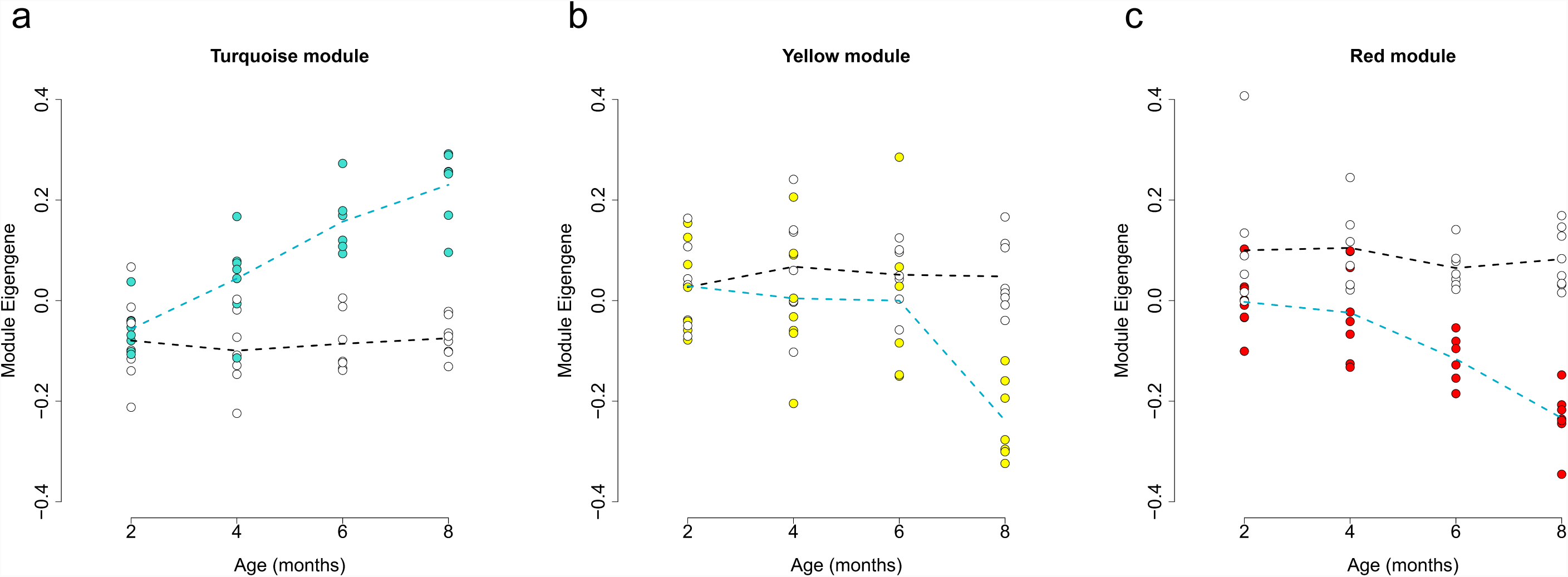
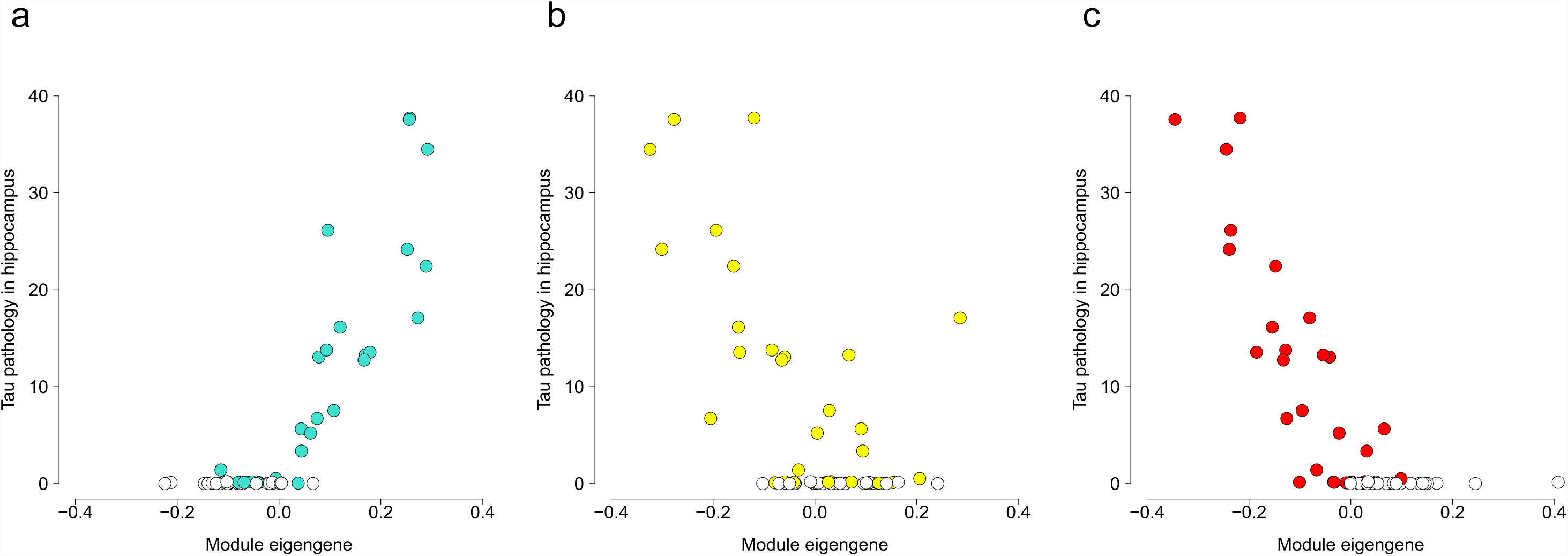
Variation in three entorhinal cortex co-expression modules parallels the accumulation of tau pathology in rTg4510 mice. Shown are module eigengene values for each individual mouse at four time-points for **(a)** the turquoise module (n = 3091 transcripts, linear regression, F(3,50) = 12.18, β = 0.28, *P* = 4.23E-06), **(b)** the yellow module (n = 1102 transcripts, linear regression, F(3,50) =5.79, β = −0.29, *P* = 0.0018), and **(c)** the red module (n = 726 transcripts, linear regression, F(3,50) = 5.58, β = −0.21, *P* = 0.0022). Total n = 59 animals (6-8 animals per group). The same three modules are correlated with levels of tau pathology quantified using immunohistochemistry in the same individuals. Shown are scatter-plots highlighting the **(d)** positive correlation between module eigengene in the turquoise module and tau pathology in the hippocampus (Pearson correlation, r = 0.85, *P* = 5.00E-17), **(e)** negative correlation between module eigengene in the yellow module and tau pathology in the hippocampus (Pearson correlation, r = −0.63, *P* = 1.00E-07), and **(f)** negative correlation between module eigengene in the red module and tau pathology in the hippocampus (Pearson correlation, r = −0.79, *P* = 2.00E-13). Coloured circles represent rTg4510 TG mice and white circles represent WT control mice. Each circle represents a single individual. Dashed lines represent mean paths for each time point. Total n = 58 animals (6-8 animals per group).

Within each of these three modules we ranked transcripts based on their intramodular connectivity to identify “hub” genes within each network, finding many genes known to play a major role in the neuro-immunological and neurodegenerative processes involved in AD. In the turquoise module the four genes with the highest intramodular connectivity (i.e. those with most connections to other genes) were *Cd63, Msn, Npc2* and *Tnfrsf1a* (**Supplementary Table 17)**, with other highly interconnected transcripts including several genes identified as having a role in LOAD from GWAS (e.g. *Abca1, Clu* and *Apoe*) in addition to genes previously implicated in AD pathology (e.g. *Itgax, Clec7a* and *Cd68*). Furthermore, genes identified as having the strongest connections (edges) to other genes (nodes) in the turquoise module included *C1qb, Mpeg1, Tyrobp*, and *Trem2* (**Figure 6a**). In the yellow module the four genes with the highest intramodular connectivity were *Atp9a, Ywhag, Rab3a* and *Svop* (**Supplementary Table 18)**, with *App* also being a highly-connected gene in this module. Genes identified as having the strongest connections to other genes in the yellow module included *Atp9a, Faim2, Ppp2r1a* (**Figure 6b**). In the red module *Atxn7l3, Sept5, Cbx6* and *Fbxl16* were the top most connected genes (**Supplementary Table 19**). Genes with the strongest connections to other genes in the red module included *Dlgap3, Shank3, Epn1*, and *Fbxl16* (**Figure 6c**).

**Figure 6.**
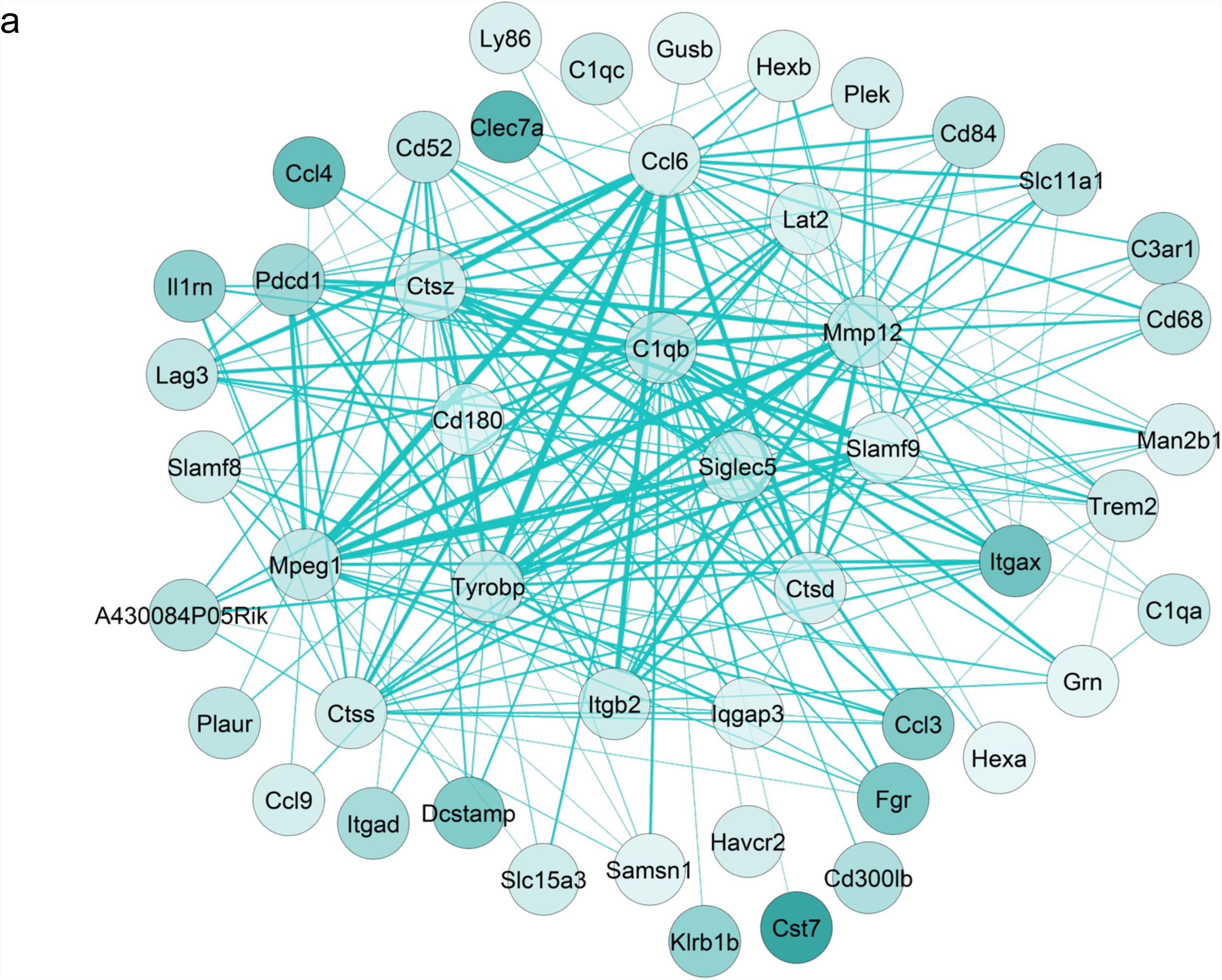
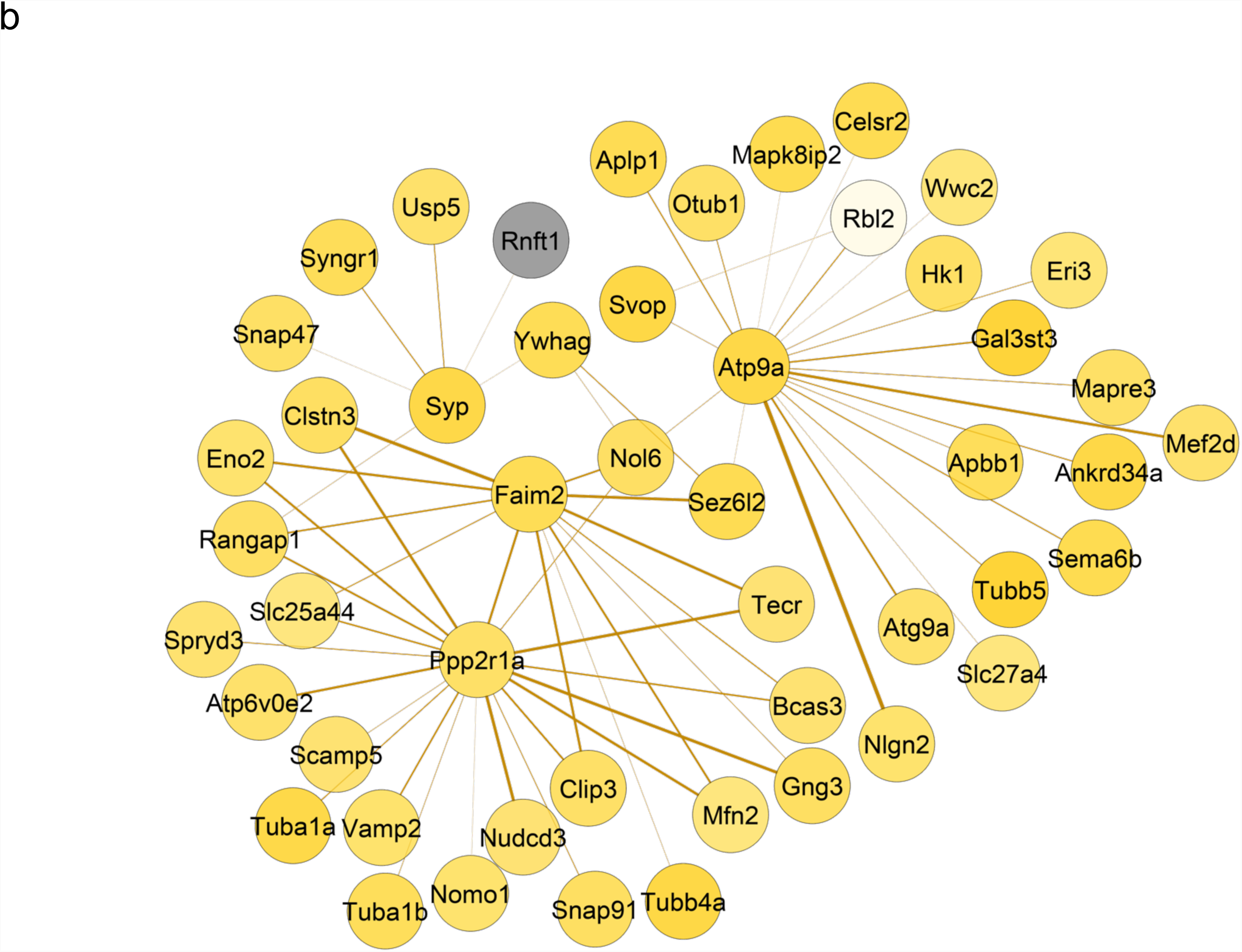
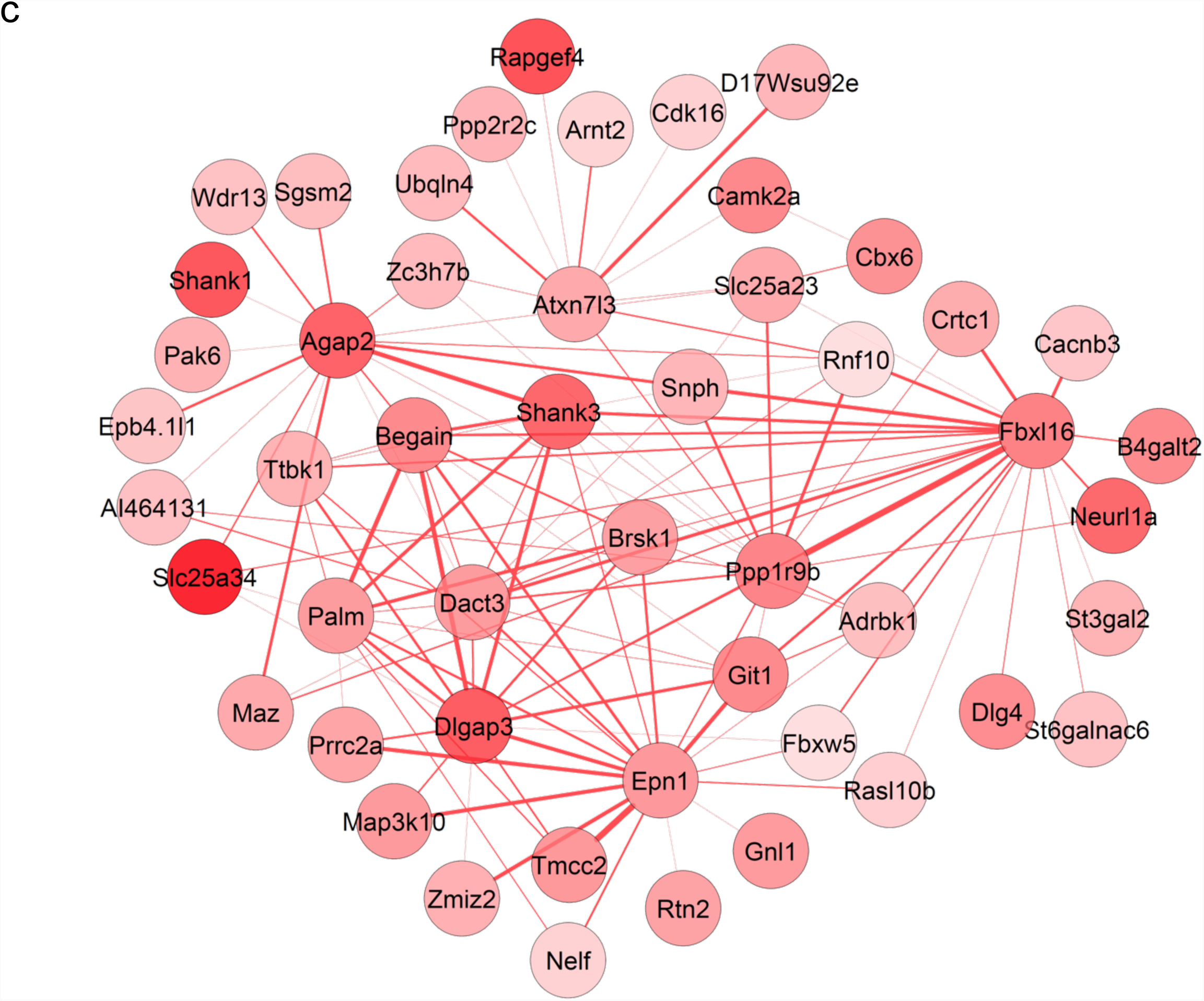
Network plots highlighting core members of gene co-expression modules associated with the development of tau pathology. Shown are the top 50 nodes (i.e. genes) with the strongest edges (representing individual connections with other genes) for each associated module. **(a)** Turquoise module (all upregulated genes – log2 fold change for latest time point against baseline). **(b)** Yellow module (downregulated genes in yellow, and upregulated genes in grey). **(c)** Red module (all downregulated genes). Stronger colors reflect higher absolute log2 fold change (8 months against 2 months). Total n = 58 animals (6-8 animals per group).

### Co-expression changes identified in rTg4510 mice overlap with AD-associated co-expression changes from human studies

We next compared the significant rTg4510 co-expression modules to AD-associated co-expression modules reported in a recent human post-mortem RNA-seq meta-analysis, focusing on modules identified in dorsolateral prefrontal cortex (DLPFC) and temporal cortex (TCX)^53^. Briefly, we used a hypergeometric test to identify overlaps between the six rTg4510-associated co-expression modules (“salmon”, “turquoise”, “purple”, “yellow”, “light-cyan”, and “red”) and four DLPFC (**Supplementary Table 20**) and five TCX (**Supplementary Table 21**) AD-associated human co-expression modules, restricting our analysis to mouse-human homologs (see **Methods**). After controlling for the number of comparisons performed for each of the human brain regions (DLPFC: P < 0.0021; TCX: P < 0.0017), each of the rTg4510-associated modules was found to significantly overlap with at least one AD-associated module in both human cortical regions. For example, genes in the turquoise rTg4510 module (enriched for pathways involved in activation of the immune system (**Supplementary Table 13**)), were found to overlap significantly with two human DLPFC modules (“DLPFC-blue” and “DLPFC-brown” from Logsdon et al.^53^) and three TCX modules (“TCX-blue”, “TCX-turquoise” and “TCX-yellow” from Logsdon et al.^53^) associated with AD; for this module the largest proportion of overlaps in genes were found with the “DLPFC-blue” module (n = 658 genes, 40.82% of the human module gene list, P < 2.2E-16) and the “TCX-turquoise” module (n = 389 genes, 39.1% of the human module gene list, P < 2.2E-16). Interestingly, GOseq analysis highlighted a strong enrichment for immune response processes amongst the rTg4510 turquoise module genes overlapping with those in both the “DLPFC-blue” module (**Supplementary Table 22**) and the “TCX-turquoise” module (**Supplementary Table 23**). Reflecting the similarities between these two human cortex modules, the list of overlapping genes includes many of the core hub transcripts identified in the turquoise rTg4510 turquoise module for both the “DLPFC-blue” module (e.g. *CD63, ABCA1, CLU, APOE, ITGAX, CLEC7A, C1QB, TYROBP*, and *TREM2*) and “TCX-turquoise” module (e.g. *CD63, ITGAX, CLEC7A, C1QB, TYROBP*, and *TREM2*). Together, these results indicate that the transcriptional networks associated with tau pathology in rTg4510 mice overlap considerably with those identified in human AD cortex and are involved in driving common molecular pathways.

## CONCLUSIONS

In this study, we identified transcriptional changes in the entorhinal cortex associated with the progression of AD-associated pathology in transgenic models of both tau (rTg4510) and amyloid (J20) pathology. We found robust genotype-associated differences in entorhinal cortex gene expression in both models and identified widespread changes in gene expression paralleling the development of tau pathology in rTg4510 mice and reflecting alterations observed in other models of tau pathology. Of note, the list of transcripts progressively altered in rTg4510 mice includes genes robustly associated with familial AD from genetic studies of human patients, including *App* which is a key driver of amyloid pathology. It also includes genes annotated to both common and rare variants identified in GWAS and exome-sequencing studies of late-onset sporadic AD. Systems-level analyses identified discrete co-expression networks associated with the progressive accumulation of tau, with these also being enriched for genes and pathways previously implicated in neuroimmune and neurodegenerative processes driving AD pathology. Further support for upregulation of immune system genes in response to tau pathology comes from our finding of increased expression of complement pathway genes including *C1qa, C1qb*, and *C1qc*. Finally, we compared these tau-associated networks to those identified in human post-mortem tissue from AD individuals, finding considerable overlap with disease-associated co-expression modules.

To our knowledge, our study represents the most systematic analysis of transcriptional variation in mouse models of tau and amyloid pathology, and is the first to focus specifically on changes in the entorhinal cortex, a key region of the brain implicated early in the pathogenesis of AD^18^. Compared to previous studies of transcriptional variation in transgenic mouse models of AD we profiled a relatively large number of samples spanning multiple time-points selected to encompass the development of pathology; our study was therefore well powered to identify gene expression differences associated with both genotype and the progression of AD pathology. Furthermore, we implemented a statistical approach that enabled us to detect progressive changes in gene expression across age between the TG and WT samples, not only identifying stable differences induced by the transgene at each time-point, but also assessing temporal transcriptional changes relative to baseline *within* mutant mice. Our detailed immunohistochemical analyses also allowed us to directly compare transcriptional variation with measures of tau and amyloid pathology measured in the same individual mice.

Despite these strengths, our study has a number of important limitations that should be considered when interpreting these results. First, to minimize the heterogeneity in our analysis we only profiled female mice. However, a number of sex differences have been previously reported for these models, with females demonstrating elevated and more progressive pathology than males^24,54^. Future work should focus on examining the extent to which the transcriptional profiles identified here are consistent between male and female mice. Second, our analysis was performed on bulk entorhinal cortex tissue, comprising a mix of different neural cell-types; consequently, changes in the fractional contribution of any given cell type to the total cellular population will contribute to the observed outcomes at each time-point. Given the compelling evidence in our data for an enrichment of microglial markers, previously shown to be upregulated in AD^42,43^, as well as upregulation of canonical markers of astrocytes, future work should focus on identifying changes that occur within these and other brain cell-types. Of note, immunocytochemistry analyses of tissue sections from the left-brain hemisphere of these mice revealed a progressive increase in the microglia/macrophage marker Iba1, indicating that our bulk-tissue RNA-seq measurements reflect real underlying cellular changes. In rTg4510 mice it is also interesting to consider neuron-specific genes that are not downregulated in what is a falling total neuronal population; these might represent transcripts that are actually upregulated in response to neuropathology in neuronal cells. Third, compared to the rTg4510 model, relatively few transcriptional changes were observed in J20 mice, potentially reflecting the slower and later accumulation of pathology^48^, as well as the potential absence of neurodegeneration, in the entorhinal cortex in this model; future work should focus on the analysis of other brain regions more directly affected in the early stages of amyloid pathology. Interestingly, however, we found that effect sizes for the transcripts identified as being progressively dysregulated in rTg4510 mice were significantly correlated across both models, suggesting some common transcriptional mechanisms are involved in both tau and amyloid pathology.

In summary, we provide compelling evidence for widespread transcriptional changes in the entorhinal cortex paralleling the progression of AD pathology. Our data suggest that the altered expression of multiple genes, including several known AD risk genes is robustly associated with the accumulation of tau, with tau-associated co-expression networks overlapping those altered in human AD cortex. Our data provide further support for an immune-response component in the accumulation of tau, and reveal novel molecular pathways associated with the progression of AD neuropathology.

## METHODS

### Mouse samples

All animal procedures were carried out at Eli Lilly and Company, in accordance with the UK Animals (Scientific Procedures) Act 1986 and with approval of the local Animal Welfare and Ethical Review Board. rTg4510 (rTg(tet-o-TauP301L)4510)^19,20^, licensed from the Mayo Clinic (Jacksonville, FL, USA), were bred on a mixed 129S6/SvEvTac + FVB/NCrl background (heterozygous tau responder x heterozygous tTA effector). Bi-transgenic (CC, here referred as TG) female mice and littermate controls (WW, here identified as WT), 2, 4, 6 and 8 months-old (n = 9-10 animals per group), were used for this study. J20 (B6.Cg-Zbtb20Tg(PDGFB-APPSwInd)20Lms/2Mmjax)^22,26^, licensed from Gladstone Institute (San Francisco, California, United States), with founder mice purchased from MMRRC at The Jackson Laboratory (Bar Harbor, Maine, United States), were bred on a C57BL/6JOlaHsd background (parental generation: hemizygous male x wild type female). Hemizygous (here identified as TG) females and littermate controls (WT), 6, 8, 10 and 12 months of age (n = 9-10 animals per group), were used for this study. All mice were bred and delivered to Eli Lilly and Company (Windlesham, UK) by Envigo (Loughborough, UK). At Eli Lilly, animals were housed under standard conditions (constant temperature and humidity) with a 12h light/dark cycle in individually ventilated cages (up to 5 animals per cage), with free access to food (Teklad irradiated global rodent diet (Envigo, United Kingdom)) and water. Mice were terminally anaesthetized with pentobarbital (intraperitoneal injection) and transcardially perfused with phosphate-buffered saline (PBS). The entorhinal cortex was dissected from the left brain hemisphere on wet ice (according to Heffner et al.^55^) and snap-frozen on dry ice for subsequent RNA-seq analysis. The right brain hemisphere was immersed in 10% buffered formalin for fixation (7-8 days) and processed for subsequent immunohistochemistry pathology assessments.

### Histopathology

The right hemisphere from all animals was processed using the Tissue TEK® VIP processor (GMI Inc) and embedded in paraffin wax. 6 μm serial sagittal sections (from bregma 0.84 to 1.08) were obtained using rotary microtomes (HM 200 from Ergostar and HM 355S from Thermo Scientific), with sections mounted on glass slides (two sections per slide). Negative and positive controls were used for each immunohistochemistry experiment. Deparaffinisation of the tissue was achieved using xylene (Fisher Scientific), followed by 70% ethanol (industrial methylated spirit, Fisher Scientific) and deionised water for rehydration of the sections. Heat induced epitope retrieval was performed in a PT Module (Thermo Scientific) containing citrate buffer (dilution 1:100). Samples were blocked using normal goat serum (Vector labs, catalogue number S-1000). To assess tau pathology, we used mouse monoclonal PG5 (provided by Peter Davies from Albert Einstein College of Medicine, Bronx, NY, USA)^56^ as the primary antibody (1:8000), which recognizes tau phosphorylated at Ser409, and biotinylated goat anti-mouse IgG (Vector labs, catalogue number BA-9200, lot number 2B0324) as the secondary antibody (1:200), as previously described^57^. To assess amyloid pathology, we used mouse monoclonal biotinylated 3D6 (b3D6, provided by Eli Lilly, 1:1000), which binds to the amino acids 1-5 in amyloid beta (Aβ)^58^. All samples for each mouse model were immunostained together in an autostainer (Autostainer 720 for PG-5 and 720N for b3D6, Thermo Scientific). For detection we undertook enzymatic labelling using peroxidase (Vectastain Elite ABC HRP Reagent, Vector Laboratories) and DAB substrate (Vector Laboratories). Images were digitised with Scanscope AT slide scanner (Aperio) at 20x magnification. Visualization of the digitized tissue sections and delineation of the regions of interest (hippocampus, cortex and thalamus) were achieved using Imagescope software (version 12.2.1.5005; Aperio). Positivity was quantified automatically using a positive pixel algorithm calibrated to ignore non-specific staining, and the burden of tau or amyloid pathology was expressed as percentage area. Statistical analysis (mixed factorial ANOVA) was performed using Microsoft Excel 2013.

### RNA isolation and sequencing

Samples were labelled with anonymized ID codes and processed in batches, blinding genotype from the experimenter/analyst for individual samples. Tissue samples from each model were processed separately and individual samples were randomized to ensure that each group was equally represented in each processing batch. Total RNA from all samples was isolated from the entorhinal cortex using the AllPrep DNA/RNA Mini Kit (Qiagen), with minor modifications to the manufactuer’s protocol. Briefly, we added lysis buffer (containing added β-mercaptoethanol) to each tissue sample, disrupted the tissue using a homogenizing pestle, and homogenized the lysate using a pipette. The lysate was centrifuged and the supernatant removed and transferred to an AllPrep DNA spin column. After centrifugation, the flow-through was used for RNA purification by mixing with 70% ethanol, running it through the RNeasy spin column (including DNase treatment) and eluting in RNase-free water. RNA quality and quantity for all samples was checked using RNA ScreenTape (Agilent) with the optimal eight samples for each group (RIN ≥ 8, Supplementary Table 1 and Supplementary Table 2) selected for transcriptional profiling (total n = 128 samples; two models (rTg4510/J20) x two groups (TG/WT) x four time-points x eight individual animals per group). Stranded-specific mRNA sequencing libraries were prepared using the TruSeq Stranded mRNA Sample Prep Kit (Illumina) using the Bravo Automated Liquid Handling Platform (Agilent). cDNA libraries were prepared from ∼450ng of total RNA plus ERCC spike-in synthetic RNA controls^59^ (Ambion, dilution 1:100). Libraries were individually cleaned up using Ampure XP magnetic beads (Beckman Coulter), their concentrations were determined using D1000 ScreenTape System (Agilent), and samples were pooled together to a 2nM concentration, for subsequent sequencing (three pools of 22 samples for J20 samples and one pool of 64 samples for rTg4510 samples). Pooled libraries were quantified using a Qubit Fluorometer (Thermo Fisher Scientific), Tapestation HS ScreenTape System (Agilent Technologies), and qPCR. Final library pools were distributed across twelve HiSeq2500 (Illumina) lanes (six lanes for each model) and subjected to 125bp paired-end sequencing yielding a mean untrimmed read depth of ∼20 million reads/sample (**Supplementary Table 1** and **Supplementary Table 2**).

### RNA-seq data processing

All sequencing data processing was performed on a Unix-based operating system server. Raw files were demultiplexed into *FASTQ* files (Phred (Q) ≥ 35, **Supplementary Table 1** and **Supplementary Table 2**) and checked for potential contamination. The randomized FASTQ files underwent quality control (QC) assessments using *FastQC*^60^ (version 0.11.4). Trimming (ribosomal sequences removal, quality threshold 20, minimum sequence length 35) was performed with f*astqmcf*^61^ (version 1.0) and trimmed samples were aligned to the *mm10* (GRCm38.p4) reference mouse genome using *STAR*^*^62^*^ (version 2.5.3a), with mapping ≥ 85% (**Supplementary Table 1** and **Supplementary Table 2**). Gene expression quantification (quantification of fragments or templates, hereby referred as read counts) was achieved using *featureCounts*^63^ (version 1.5.2). Following confirmation of genotype and QC, 7 samples were excluded from subsequent analysis leaving a final number of 121 high-quality RNA-seq datasets (6-8 animals per group).

### Gene expression analysis

All analyses were performed in R (version 3.4.3) unless otherwise stated. Read counts were analysed for differential expression using the R package *DESeq2*^*64*^ (version 1.16.1) downloaded from Bioconductor^14^.DESeq2 uses the raw read counts, applies an internal normalization method, and does estimation of library size, estimation of dispersion, and negative binomial generalized linear model fitting^64^. Data sets were filtered for non-expressed and lowly expressed genes (minimum of 6 counts across all samples), and similarity in the genome-wide expression profile between samples was visualized in a heatmap clustered by Euclidean distance (**Supplementary Figure 21 and Supplementary Figure 22**) and a principal component analysis (PCA) plot of the first two principal components (**Supplementary Figure 23** and **Supplementary Figure 24**). We were interested in detecting both genotype effects and progressive changes across age between the transgenic and wild type samples. We used the following statistical model, including main effects for both Genotype and Age (both coded as categorical variables) and an interaction between these two terms:

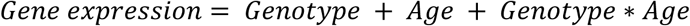

To identify significant Genotype effects a Wald test was used, and to identify significant effects of Age and interaction effects (i.e. Genotype*Age) we used the likelihood-ratio test, both applied with the *DESeq* function from the *DESeq2* package^64^. *P* values were adjusted for multiple testing, using the false discovery rate (FDR) method (also known as Benjamini and Hochberg correction)^65^ implemented with the R function *p.adjust*; FDR-adjusted *P* values < 0.05 were defined as significant. Potential differences in proportions of upregulated versus downregulated genes as well as overlapping fold changes in both models were interrogated using the binomial test. Functional annotation and gene ontology analyses were done with *GOseq^66^* (1.30.0), based on genes with FDR < 0.05.

### Quantifying human transgene expression

Mouse and human *App*/*APP* and *Mapt*/*MAPT* sequences were compared using BLAT^67^ for divergent transcript sequences representing specific mouse and human gene sequences. Two 200bp regions spanning 4 exons were chosen as representative mouse-specific *App*. Similar regions consisting of two 200bp exonic regions were also chosen for human *APP*. Mouse-specific *Mapt* and human-specific *MAPT* sequences were chosen from a 2kb region present in the 3’UTR. Using *Bowtie2*^*68*^ (version 2.3.4.3), indices based on these sequences were then built, and alignments were performed using the *FASTQ* read 1 sequences. Counts of read alignments for mouse and human specific indices were then plotted as a ratio of unique (mouse or human) reads relative to the total number of input reads.

### Comparison with RNA-seq data from the Mouseac database

RNA-seq data (transcripts per million, TPM) from two mouse models^14,51^ (TAU (CaMKII-MAPTP301L) and TAS10 (SwAPP, K670N/M671L)) were downloaded from the Mouseac online database (www.mouseac.org), with corresponding detailed phenotypic data downloaded from GEO^28,29^ (accession number GSE64398). Only genes identified as differentially expressed (FDR < 0.05) in our analysis (for rTg4510 and J20 mice) were kept for further statistical analysis. TPM was log transformed (log2(x+1)) and the same linear regression model described above (*Gene expression* = *Genotype* + *Age* + *Genotype* * *Age*), using ANOVA to test for significant differences associated with either the Genotype, Age, or Genotype*Age terms, was used. *P* values were corrected for the number of genes compared across datasets using Bonferroni correction. Potential differences in proportions of upregulated versus downregulated genes as well as overlapping fold changes in both models were interrogated using the binomial test.

### Co-expression network analysis

Using weighted co-expression network analysis^69,70^ (WGCNA) (version 1.63), we constructed a signed co-expression network for each mouse model using log transformed counts from all samples. Logarithmic transformation of raw counts was achieved using the rlog function from DESeq2, which minimizes variability in genes with low counts^64^. We checked the data for missing values and outliers, and removed one sample from the analysis for the rTg4510 dataset (flagged as an outlier) before building the networks. Signed WGCNA co-expression networks were built using the lowest power for which the scale-free topology fit index curve flattened out after reaching 0.90 resulting in a soft-threshold power of 10 and 9 for rTg4510 and J20 datasets, respectively, and a minimum module size of 30. For each module of highly interconnected genes, colour-labelled according to the WGCNA conventions, we calculated the module eigengenes (MEs) as the first principal component of the expression matrix, which provide a representative expression profile for each module^69^. In order to identify modules significantly associated with pathology burden, we calculated correlation coefficients between these MEs with the available pathology data and explored the most significant associations. In addition we used the same linear regression model as described for the gene-level analysis (*ME* = *Genotype* + *Age* + *Genotype* * *Age*) using ANOVA to test for significant differences due to either the Genotype, Age, or Genotype*Age terms. *P* values were corrected for multiple comparisons using the Bonferroni correction, to correct for 18 modules in the rTg4510 (statistical threshold was adjusted to 0.05/18 = 0.0028) and 21 modules in the J20 (statistical threshold was adjusted to 0.05/21 = 0.0024) mice. We used the *GOseq* R package (version 1.30.0) to perform functional annotation and gene ontology (GO) analyses for each module, where significant pathways were selected using an FDR threshold of 0.05 as previously described^66^. We used *Cytoscape*^*^71^*^ (version 3.7.0) for network visualization using the topological overlap matrix for the log transformed expression data.

### Comparison with human co-expression networks

The six rTg4510-associated co-expression modules identified in this study (“salmon”, “turquoise”, “purple”, “yellow”, “light-cyan”, and “red”), and AD-associated human co-expression modules in the dorsolateral prefrontal cortex (DLPFC) and temporal cortex (TCX)^53^, were reduced to contain only mouse-human homologs as defined by Ensembl^72^ (accessed on 14/11/2018). The level of overlap between gene members of each pair of modules was assessed via a hypergeometric test using the R (version 3.5.1) function *phyper*. *P* values were corrected for multiple comparisons using Bonferroni correction in a tissue-specific manner, where only the set of raw *P* values related to DLPFC (statistical threshold was adjusted to 0.05/24 = 0.0021) or TCX (statistical threshold was adjusted to 0.05/30 = 0.0017) modules’ overlap were considered. Using *GOseq* (version 1.30.0), we performed functional annotation and GO analyses for the common genes in each overlapping pair of modules.

## DATA AVAILABILITY

Raw RNA-seq data has been deposited in GEO under accession number GSE125957.

### ACKNOWLEDGEMENTS

This work was funded in part through the MRC Proximity to Discovery: Industry Engagement Fund (Precision Medicine Exeter Innovation Platform (PMEI Platform) ref. MC_PC_14127). IC’s doctoral studentship is supported by the Alzheimer’s Society in partnership with the Garfield Weston Foundation (grant reference 231). Data were produced by the Sequencing Service and Computational core facilities at the University of Exeter with generous support from Medical Research Council Clinical Infrastructure award (MR/M008924/1), Wellcome Trust Institutional Strategic Support Fund (WT097835MF), Wellcome Trust Multi User Equipment Award (WT101650MA) and BBSRC LOLA award (BB/K003240/1). We would like to thank Mark Ward, Katherine Sung, Alice Fisher, and Claire Cella for technical assistance with the histopathology, and Audrey Farbos for technical assistance with the RNA sequencing laboratory experiments.

## AUTHOR CONTRIBUTIONS

I.C. conducted RNA-seq and immunohistochemistry laboratory experiments. K.M. supervised RNA-seq experiments. M.J.O’N, T.K.M and Z.A. supervised animal work and histopathology experimental procedures. J.M. and A.R. obtained funding. J.M., M.J.O’N. and D.A.C. designed the study. I.C. undertook primary data analyses and bioinformatics, with analytical and computational input from E.H., A.J., E.W., E.L., H.B., J.H., and P.O’N. K.L. and A.R. helped interpret the results. I.C. and J.M. drafted the manuscript. All of the authors read and approved the final submission.

