## Supplementary Figures for "Transcriptional signatures of progressive neuropathology in transgenic tau and amyloid mouse models"

**Supplementary Figure 1. An overview of the experimental approach used in this study. (a)** To investigate transcriptional signatures of tau pathology we used the rTg4510 transgenic mouse line, which overexpresses a human mutant (P301L) form of the microtubule-associated protein tau (MAPT). To investigate amyloid pathology, we used the J20 transgenic mouse line, which expresses a mutant (K670N/M671L and V717F) form of the human amyloid precursor protein (APP). Tissue was collected from transgenic (TG) and wild type (WT) control mice at four time points. **(b)** From each mouse, the entorhinal cortex (ECX) was dissected from the left hemisphere and used for RNA isolation and subsequent RNA-seq analysis. The right hemisphere was fixed and used to quantify the progression of neuropathology across multiple brain regions using immunohistochemistry. **(c)** Our analyses focused primarily on i) identifying transcriptional differences associated with genotype and ii) identifying temporal changes in gene expression in TG mice paralleling the development of neuropathology (i.e. interactions between genotype and age).

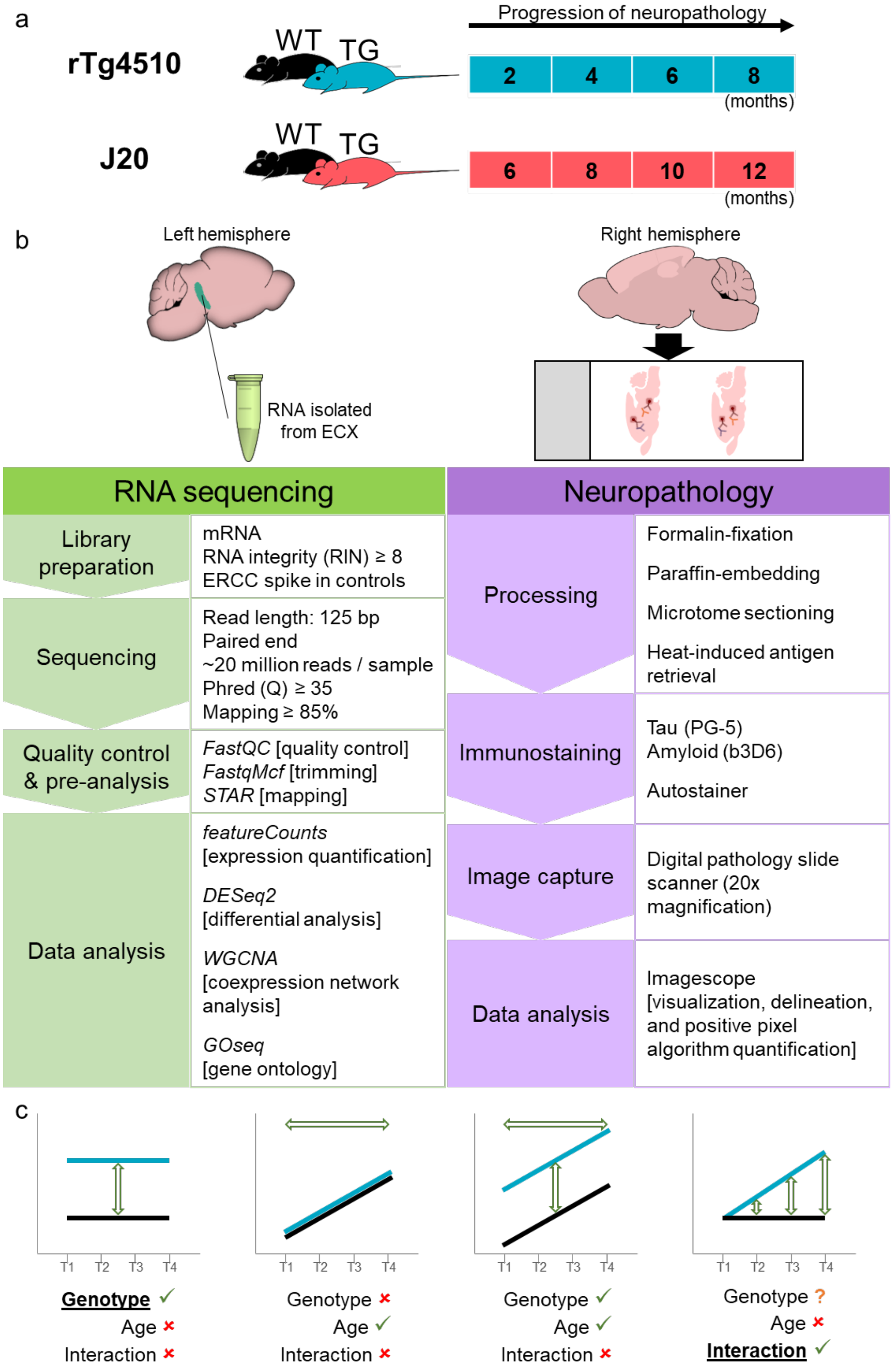

**Supplementary Figure 2. rTg4510 mice develop progressive tau neuropathology in the hippocampus and cortex. (a)** The anatomical regions of the brain tested for tau pathology: whole hippocampus (purple line), CA1 subregion of the hippocampus (light green area within the purple line), CA3 subregion of the hippocampus (brown area within the purple line), dentate gyrus (DG) subregion of the hippocampus (light red area within the purple line), whole cortex (orange line), secondary motor cortex (M2, dark yellow area within the orange line), primary motor cortex (M1, magenta area within the orange line), visual cortex (dark green area within the orange line), retrosplenial cortex (dark blue are within the orange line), thalamus (dark grey square). **(b-j)** Quantification of PG-5 immunoreactivity in each tested region: **(b)** CA1 (total n = 74 animals, 9-10 animals per group, factorial ANOVA, F(3,67) = 68.86, *P* = 1.96E-20), **(c)** CA3 (total n = 73 animals, 8-10 animals per group, factorial ANOVA, F(3,66) = 64.22, *P* = 1.51E-19), **(d)** DG (total n = 73 animals, 8-10 animals per group, factorial ANOVA, F(3,66) = 67.42, *P* = 4.58E-20), **(e)** whole cortex (total n = 72 animals, 7-10 animals per group, factorial ANOVA, F(3,65) = 53.33, *P* = 1.65E-17), **(f)** M2 (total n = 74 animals, 8-10 animals per group, factorial ANOVA, F(3,67) = 44.90, *P* = 5.01E-16), **(g)** M1 (total n = 73 animals, 8-10 animals per group, factorial ANOVA, F(3,66) = 29.28, *P* = 3.74E-12), **(h)** visual cortex (total n = 73 animals, 8-10 animals per group, factorial ANOVA, F(3,66) = 55.22, *P* = 5.64E-18), **(i)** retrosplenial cortex (total n = 73 animals, 8-10 animals per group, factorial ANOVA, F(3,66) = 38.38, P = 1.79E-14). **(j)** As expected, the thalamus, which is relatively protected from aggressive tau pathology, was characterized by relatively low levels of tau. Despite this, TG mice demonstrated significant differences in late stages (total n = 74 animals, 8-10 animals per group, factorial ANOVA, F(3,67) = 111.48, *P* = 5.41E-26). rTg4510 transgenic (TG, blue) female mice compared to wild type (WT, black) littermate control mice.

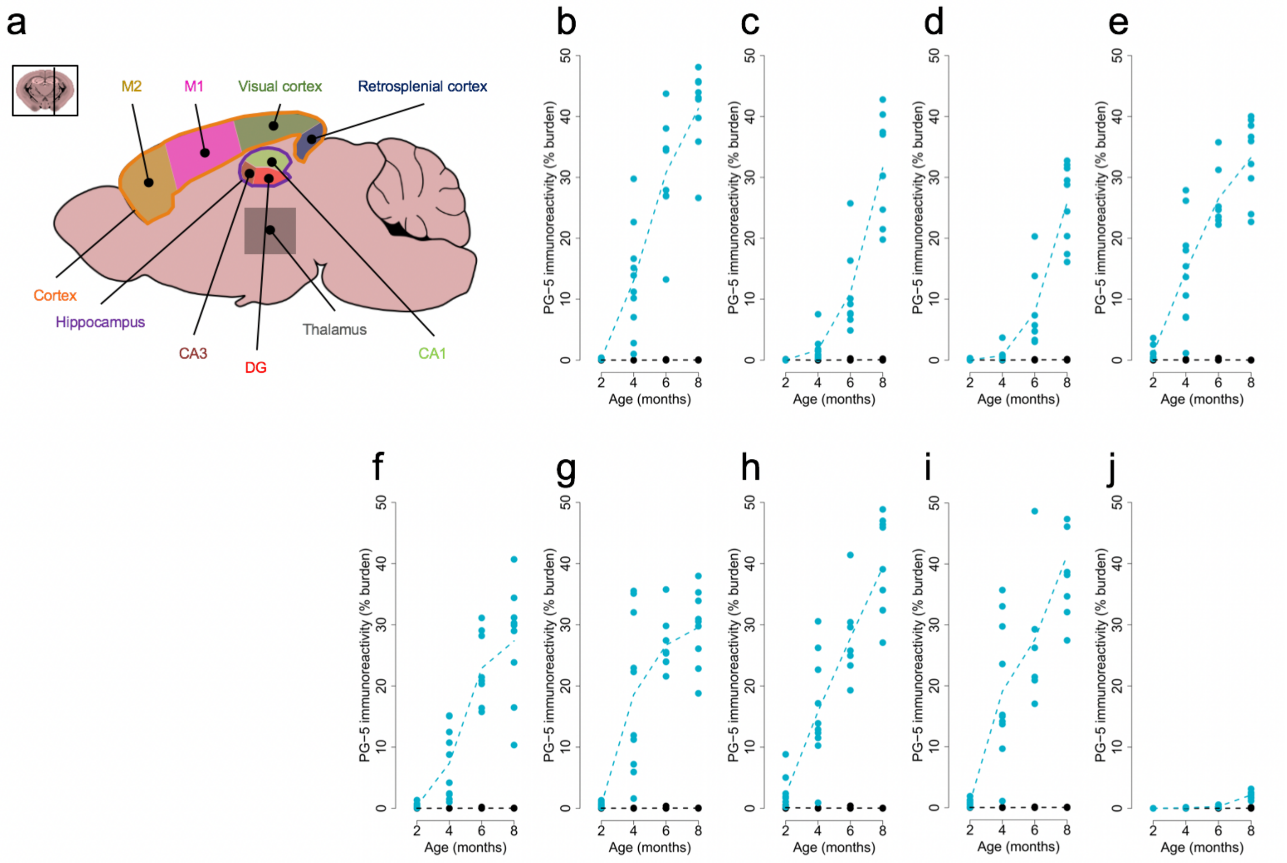

**Supplementary Figure 3. J20 mice are characterized by progressive amyloid neuropathology. (a)** Brain regions tested: whole hippocampus (purple area), whole cortex (orange line), rostral cortex (magenta area within the orange line), caudal cortex (green area within the orange line), thalamus (dark grey square). **(b-e)** Quantification of b3D6 immunoreactivity in each tested region: **(b)** whole cortex (total n = 73 animals, 8-10 animals per group, factorial ANOVA, F(3,66) = 24.75, *P* = 7.63E-11), **(c)** rostral cortex (total n = 73 animals, 8-10 animals per group, factorial ANOVA, F(3,66) = 10.63, *P* = 8.65E-06), **(d)** caudal cortex (total n = 74 animals, 8-10 animals per group, factorial ANOVA, F(3,67) = 28.62, *P* = 5.02E-12), **(e)** thalamus (total n = 73 animals, 8-10 animals per group, factorial ANOVA, F(3,66) = 1.58, *P* = 0.20). J20 transgenic (TG, red) female mice compared to wild type (WT, black) littermate control mice.

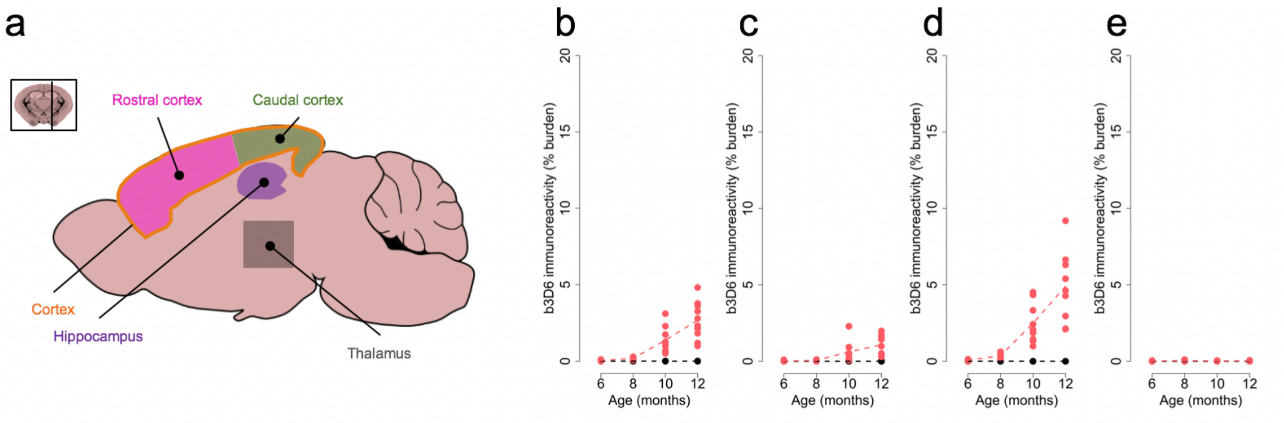

**Supplementary Figure 4. Comparison of RNA-seq reads in transgenic and wild type animals for both models.** The number of raw RNA-seq reads did not differ between WT and TG mice for comparisons of either the **(a)** rTg4510 (n = 59 animals, two-tailed unpaired t-test, t(57)= 1.35, *P* = 0.18) or **(b)** J20 (n = 62 animals, two-tailed unpaired t-test, t(60) = 0.41, *P* = 0.18) models. The proportion of RNA-seq reads uniquely mapped to the mouse genome did not differ between WT and TG mice for comparisons of either the **(c)** rTg4510 (n = 59 animals, two-tailed unpaired t-test, t(57) = -1.36, *P* = 0.089) or **(d)** J20 (n = 62 animals, two-tailed unpaired t-test, t(60) = 0.79, *P* = 0.22) models.

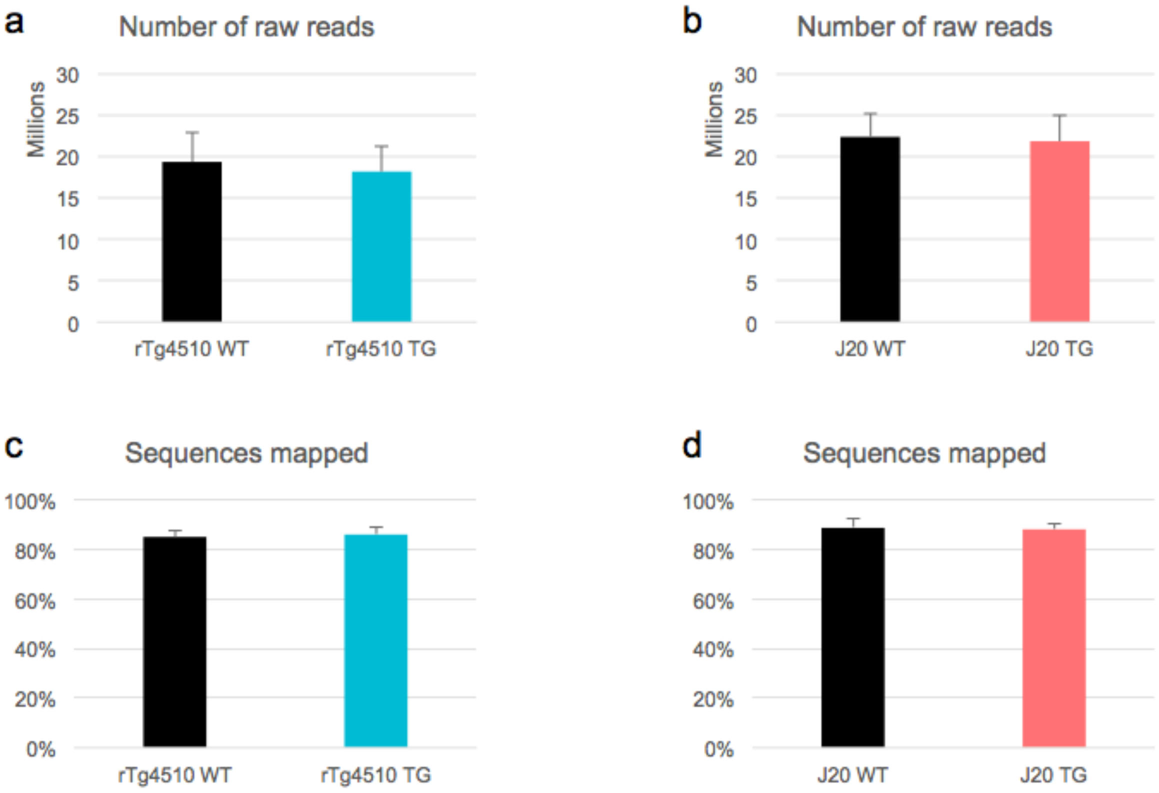

**Supplementary Figure 5. Apparent up-regulation of *Mapt* in rTg4510 mice and *App* in J20 mice results from human-specific sequences only present in transgenic mice. (a-c)** Apparent up-regulation of *Mapt* in rTg4510 transgenic (TG, blue) female mice compared to wild type (WT, black) littermate control mice (n = 59 animals). **(a)** *Mapt* gene expression from *DESeq2* highlights elevated expression of *Mapt* in TG mice (Wald test, Wald statistic = 11.11, log2 fold change = 0.50, FDR = 7.08E-25). **(b)** Ratio of unique reads that mapped to mouse-specific sections of *Mapt* or **(c)** human-specific sections of *MAPT*. **(d-f)** Apparent up-regulation of *App* in J20 transgenic (TG, red) female mice compared to wild type (WT, black) littermate control mice (n = 62 animals). **(d)** *App* gene expression from *DESeq2* highlights elevated expression of *App* in TG mice (Wald test, Wald statistic = 8.55, log2 fold change = 0.66, FDR = 2.37E-13). (e) Ratio of unique reads that mapped to mouse-specific sections of *App* or **(c)** human-specific sections of *APP*. Normalised counts were obtained using *DESeq2*. Dashed lines represent mean paths across age groups.

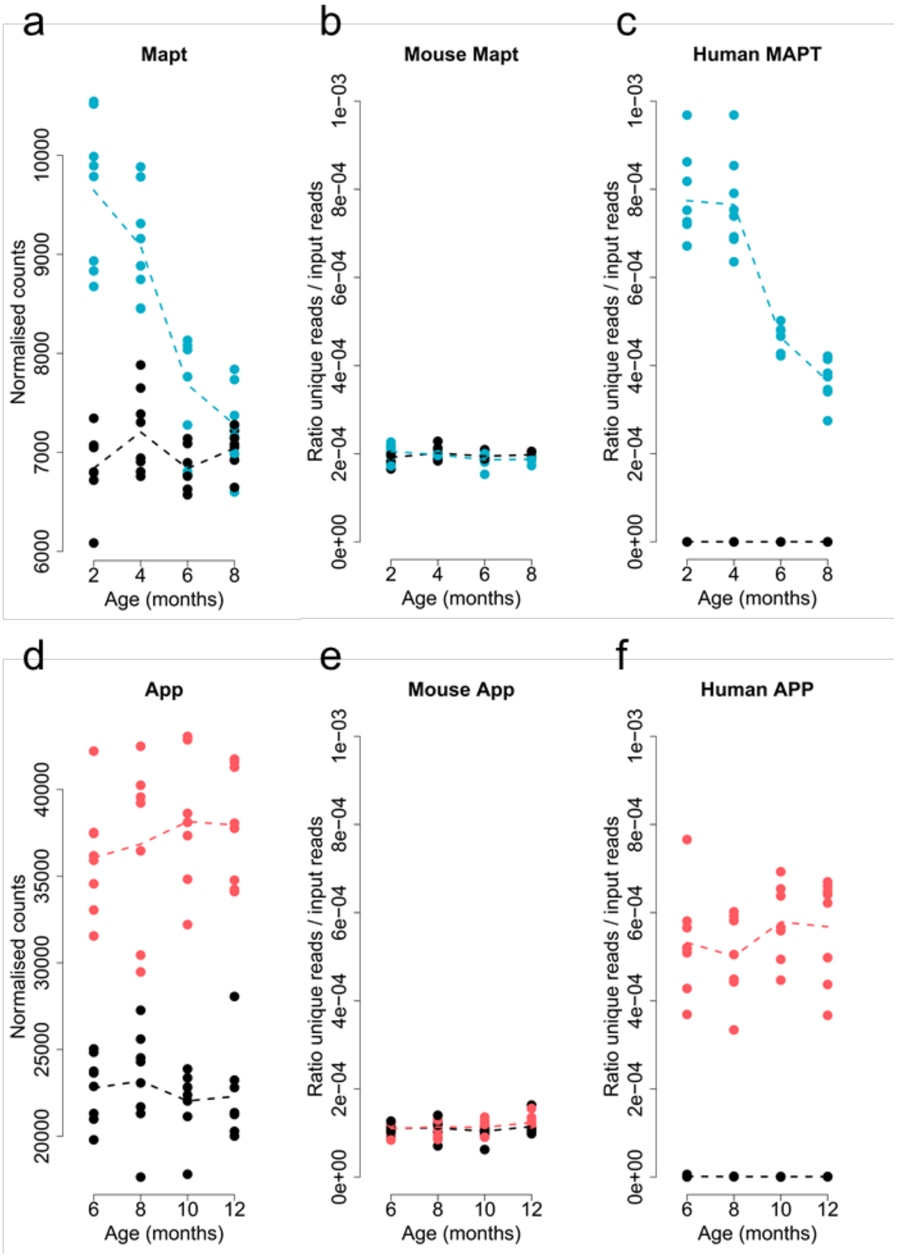

**Supplementary Figure 6. Top-ranked differentially expressed transcripts associated with rTg4510 genotype. (a)** *Car4* (Wald test, Wald statistic = 8.36, log2 fold change = 1.11, FDR = 2.41E-13). **(b)** *Gpr17* (Wald test, Wald statistic = -6.73, log2 fold change = -0.62, FDR = 5.11E-08). **(c)** *Blnk* (Wald test, Wald statistic = 6.48, log2 fold change = 0.80, FDR = 2.12E-07). **(d)** *Hspa5* (Wald test, Wald statistic = -6.16, log2 fold change = -0.58, FDR = 1.37E-06). Total n = 59 animals (29 TG, 30 WT). Normalized counts were obtained using *DESeq2*. Dashed lines represent mean paths across age groups. rTg4510 transgenic (TG, blue) female mice compared to wild type (WT, black) littermate control mice.

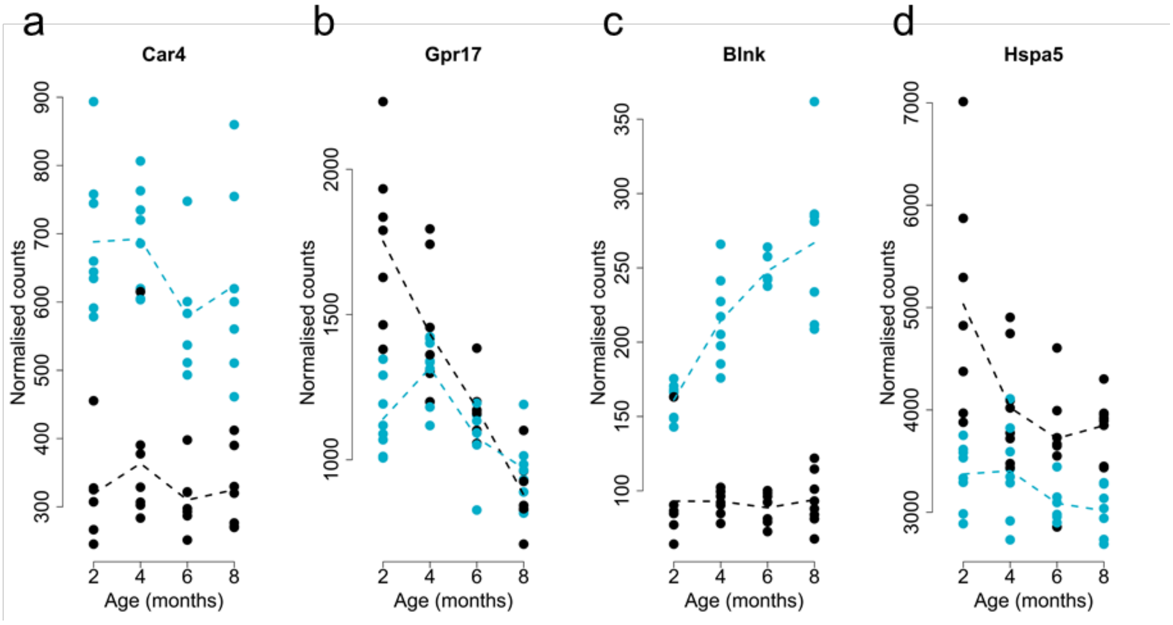

**Supplementary Figure 7. Top-ranked differentially expressed transcripts associated with J20 genotype. (a)** *Ccdc80* (Wald test, Wald statistic = 6.37, log2 fold change = 0.81, FDR = 1.74E-06). **(b)** *Abca8a* (Wald test, Wald statistic = -4.67, log2 fold change = -0.81, FDR = 0.02). **(c)** *Htr1a* (Wald test, Wald statistic = -4.48, log2 fold change = -0.51, FDR = 0.035). **(d)** *Hspa5* (Wald test, Wald statistic = -4.36, log2 fold change = -0.28, FDR = 0.049). Total n = 62 animals (30 TG, 32 WT). Normalized counts were obtained using *DESeq2*. Dashed lines represent mean paths across age groups. J20 transgenic (TG, red) female mice compared to wild type (WT, black) littermate control mice.

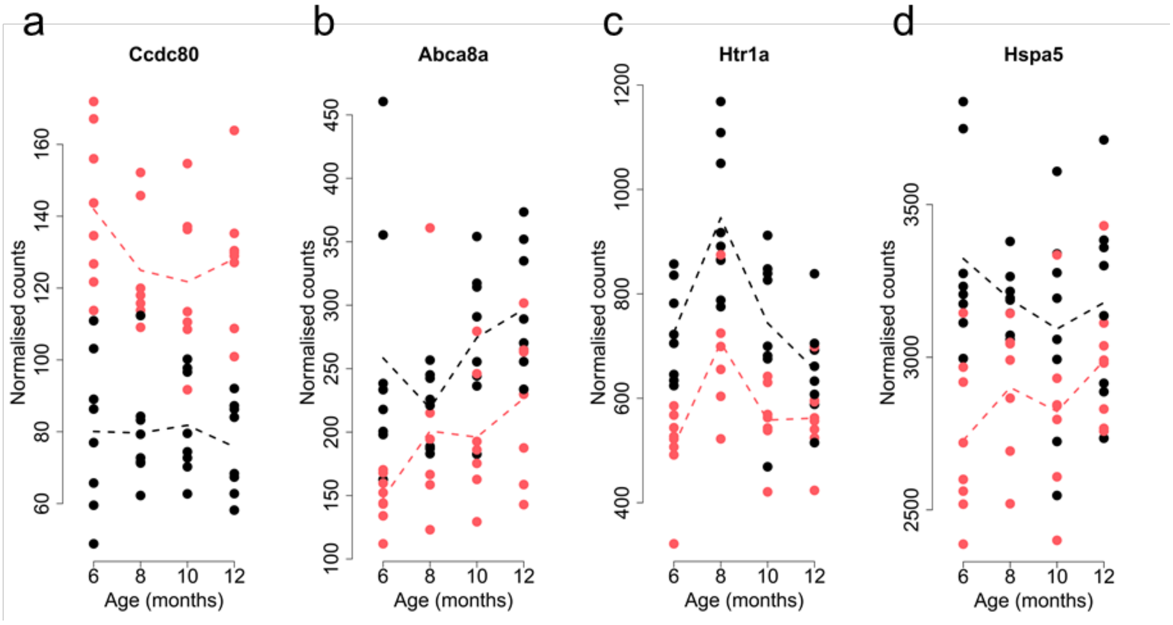

**Supplementary Figure 8. There is no overall consistency between models in effect sizes for differentially-expressed transcripts identified in each individual model. (a)** Negligible correlation for effect size (Log2 fold change) across models for differentially expressed transcripts (FDR < 0.05) associated with rTg4510 genotype (Pearson correlation, r = 0.18, P = 0.029; exact binomial test, n = 147 transcripts, P = 0.69). **(b)** No correlation for effect size (Log2 fold change) across models for differentially expressed transcripts (FDR < 0.05) associated with J20 genotype (Pearson correlation, r = 0.66, P = 0.23; exact binomial test, n = 4 transcripts, P = 0.13). Despite the overall lack of consistency between models, one transcript (*Hspa5*) is differentially downregulated (FDR < 0.05) in the same direction in both models (Tg4510: Wald statistic = -6.16, log2 fold change = -0.58, FDR = 1.37E-06; J20: Wald statistic = -4.36, log2 fold change = -0.28, FDR = 0.049).

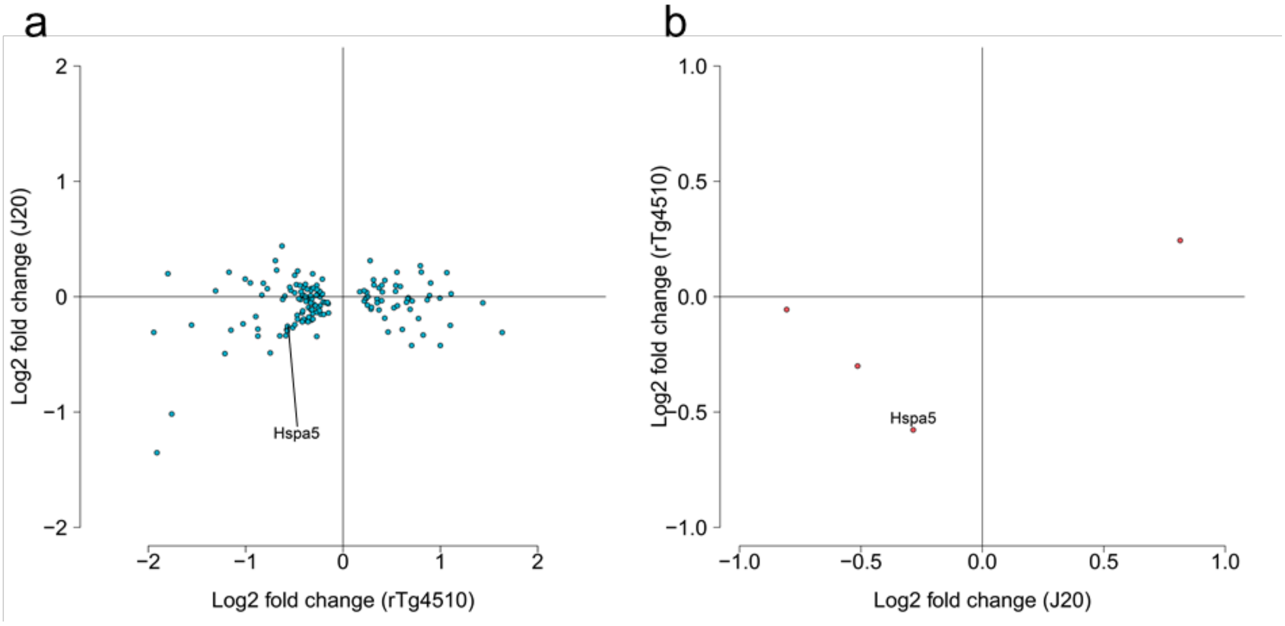

**Supplementary Figure 9. rTg4510 mice show a progressive increase in Iba1 immunoreactivity. (a-c)** Quantification of Iba1 immunoreactivity in each tested region: **(a)** hippocampus (total n = 70 animals, 7-10 animals per group, factorial ANOVA, $F(3,62) = 12.60$, $P = 1.56E-06$), **(b)** cortex (total n = 70 animals, 7-10 animals per group, factorial ANOVA, $F(3,62) = 18.13$, $P = 1.47E-08$), **(c)** thalamus (total n = 70 animals, 7-10 animals per group, factorial ANOVA, $F(3,62) = 18.85$, $P = 8.37E-09$). rTg4510 transgenic (TG, blue) female mice compared to wild type (WT, black) littermate control mice.

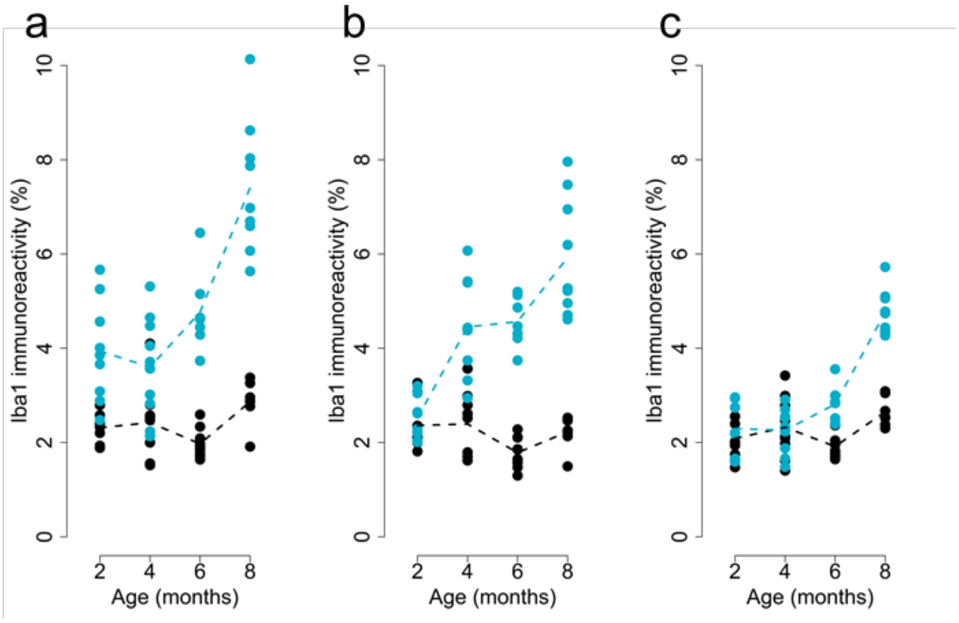

**Supplementary Figure 10. Transcripts progressively altered in rTg4510 mice include many genes implicated in familial and sporadic AD from genetic association studies.** Genes showing temporal shifts in expression in rTg4510 mice included: **(a)** *App* (Likelihood-ratio test, LRT statistic = 13.88, log2 fold change (2-8 months) = -0.35, FDR = 0.037); **(b)** *Trem2* (Likelihood-ratio test, LRT statistic = 43.82, log2 fold change (2-8 months) = 1.46, FDR = 3.73E-07); **(c)** *Pld3* (Likelihood-ratio test, LRT statistic = 36.80, log2 fold change (2-8 months) = -0.58, FDR = 5.80E-06); **(d)** *Frmd4a* (Likelihood-ratio test, LRT statistic = 27.81, log2 fold change (2-8 months) = 0.36, FDR = 0.00022); **(e)** *Clu* (Likelihood-ratio test, LRT statistic = 27.73, log2 fold change (2-8 months) = 0.80, FDR = 0.00023); **(f)** *Apoe* (Likelihood-ratio test, LRT statistic = 22.99, log2 fold change (2-8 months) = 0.86, FDR = 0.0014); **(g)** *Picalm* (Likelihood-ratio test, LRT statistic = 21.37, log2 fold change (2-8 months) = 0.26, FDR = 0.0025); **(h)** *Cd33* (Likelihood-ratio test, LRT statistic = 27.32, log2 fold change (2-8 months) = 1.14, FDR = 0.00026); and **(i)** *Abi3* (Likelihood-ratio test, LRT statistic = 17.10, log2 fold change (2-8 months) = 0.70, FDR = 0.012). Total n = 59 animals (29 TG, 30 WT). Normalised counts were obtained using *DESeq2*. Dashed lines represent mean paths across age groups. rTg4510 transgenic (TG, blue) mice compared to wild type (WT, black) littermate controls.

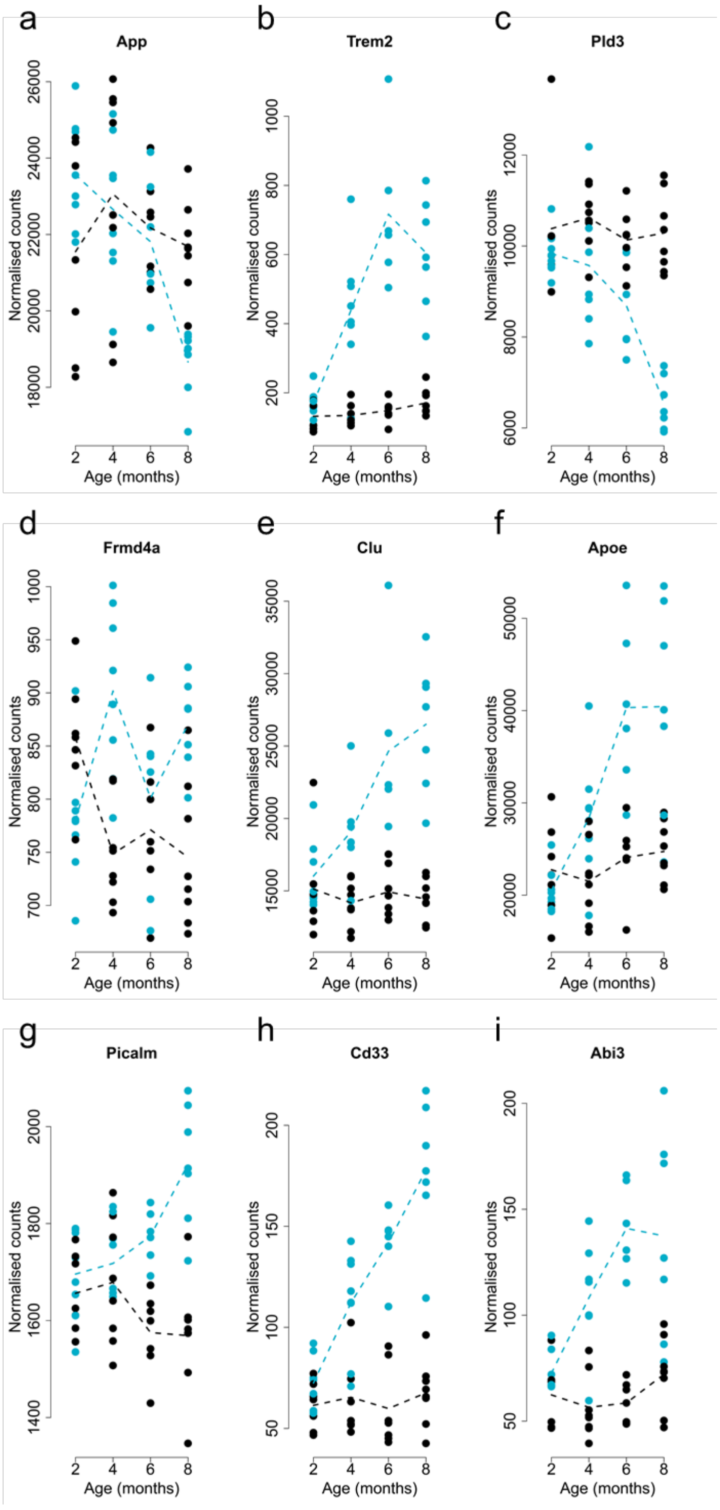

**Supplementary Figure 11. *Itgax* is upregulated with age and the progression of amyloid pathology in J20 mice.** The expression of *Itgax* in J20 mice (Total n = 62 animals, 6-8 animals per group, Likelihood-ratio test, LRT statistic = 37.37, log2 fold change (6-12 months) = 2.42, FDR = 0.00072) reflects the changes seen in rTg4510 mice (see **Figure 3d**). Normalised counts were obtained using *DESeq2*. Dashed lines represent mean paths for each time point. J20 transgenic (TG, red) female mice compared to wild type (WT, black) littermate control mice.

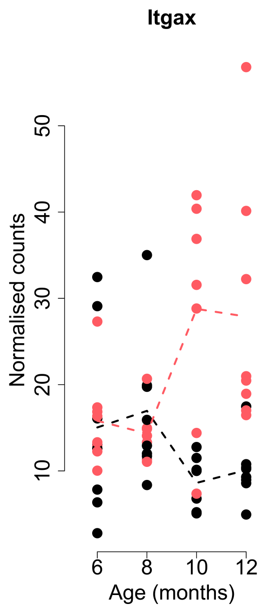

**Supplementary Figure 12. Comparison of rTg4510 genotype-associated transcriptional differences in another transgenic model of tau pathology. (a)** Correlation of effect size (log2 fold change) between rTg4510 genotype-associated genes and the same genes in the TAU mouse model from Mouseac (Pearson correlation, r = 0.33, *P* = 7.73E-5, exact binomial test, n = 138 transcripts, P = 0.11). **(b)** Correlation of genotype-associated *P* values (applying the same linear regression model) between rTg4510 and the same genes in the TAU mouse model from Mouseac (Pearson correlation, r = 0.02, *P* = 0.82).

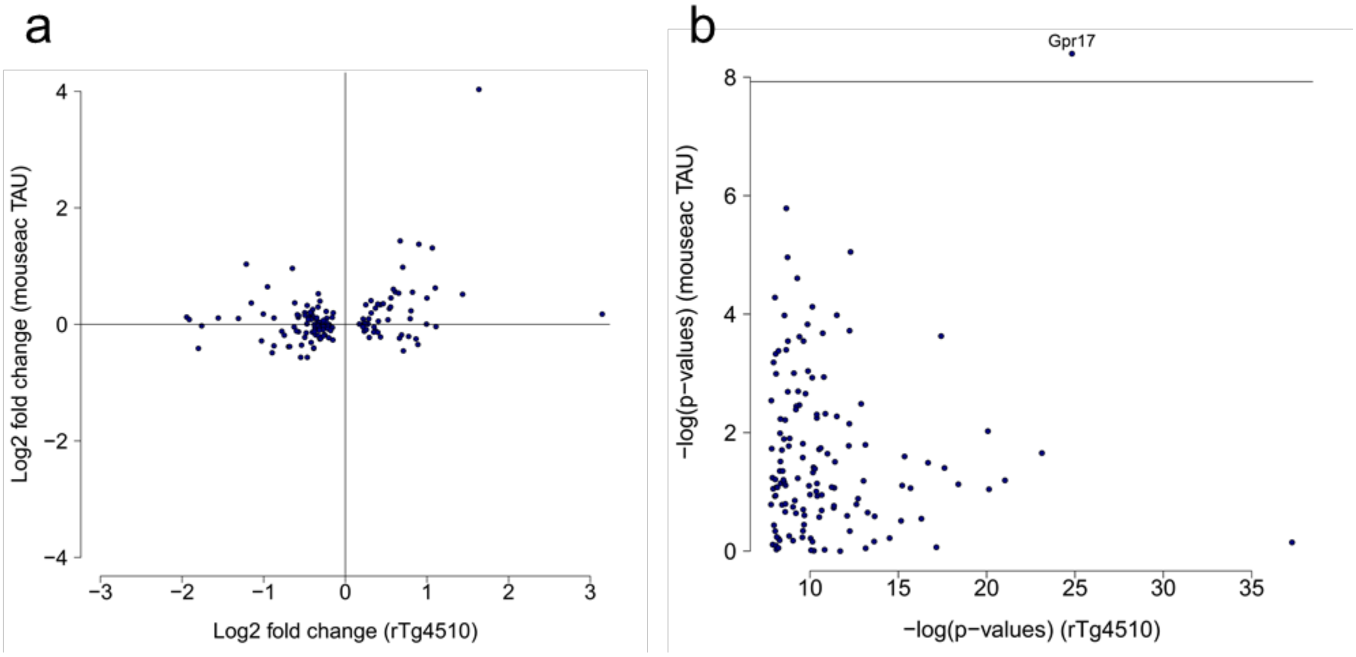

**Supplementary Figure 13. Most of top-ranked differentially expressed transcripts associated with rTg4510 genotype were not statistically replicated in another mouse model of tau pathology. (a)** *Car4* (linear regression, t(20) = -0.18, β = -1.31, P = 0.87). **(b)** *Gpr17* (linear regression, t(20) = 4.49, β = 3.13, P = 0.00023). **(c)** *Blnk* (linear regression, t(20) = 1.35, β = 1.11, P = 0.19). **(d)** *Hspa5* (linear regression, t(20) = 1.06, β = 20.72, P = 0.30). Total n = 25 animals (2-4 animals per group). TAU transgenic transcriptomic data obtained from Mouseac (dark blue) compared to wild type (black) controls. Dashed lines represent mean paths across age groups.

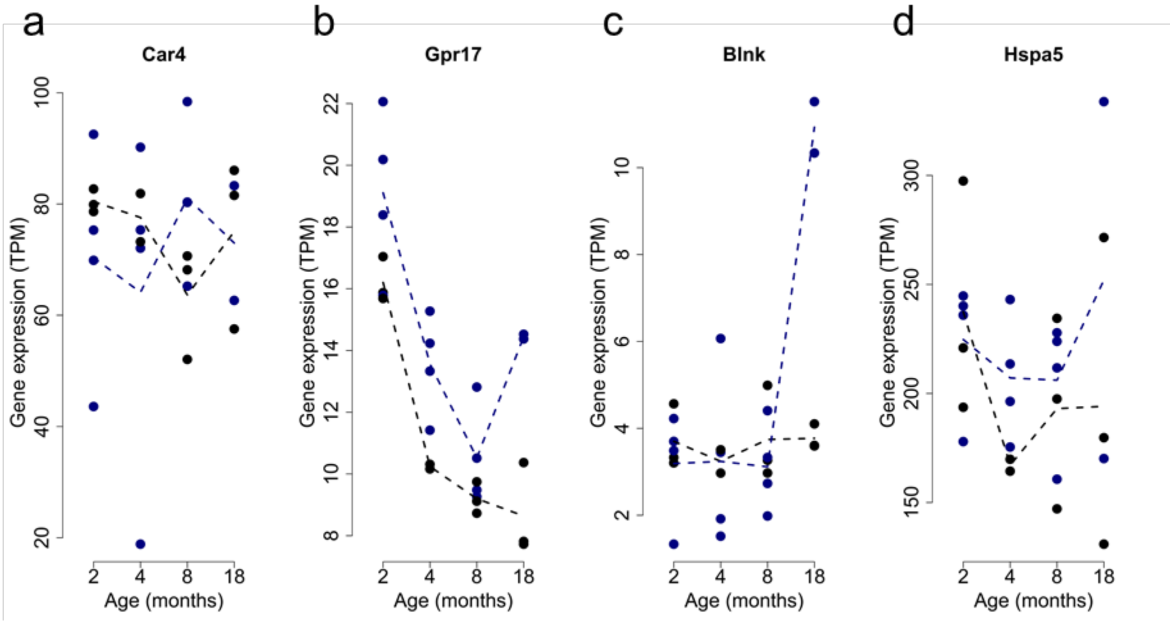

**Supplementary Figure 14. Analysis of genes showing progressive transcriptional differences in rTg4510 mice in other transgenic models. (a)** Correlation of *P* values between the rTg4510 and TAU mouse models (using transcriptional data from Mouseac) for an analysis of interactions between and genotype and age for genes identified as significant in rTg4510 mice (Pearson correlation, r = 0.46, *P* = 1.22E-86, exact binomial test, n = 1640 transcripts, *P* < 2.2e-16). **(b)** Correlation of *P* values between the rTg4510 and TAS10 mouse models (using transcriptional data from Mouseac) for an analysis of interactions between genotype and age for genes identified as significant in rTg4510 mice (Pearson correlation, r = 0.23, *P* = 3.89E-21, exact binomial test, n = 1640 transcripts, *P* < 2.2e-16).

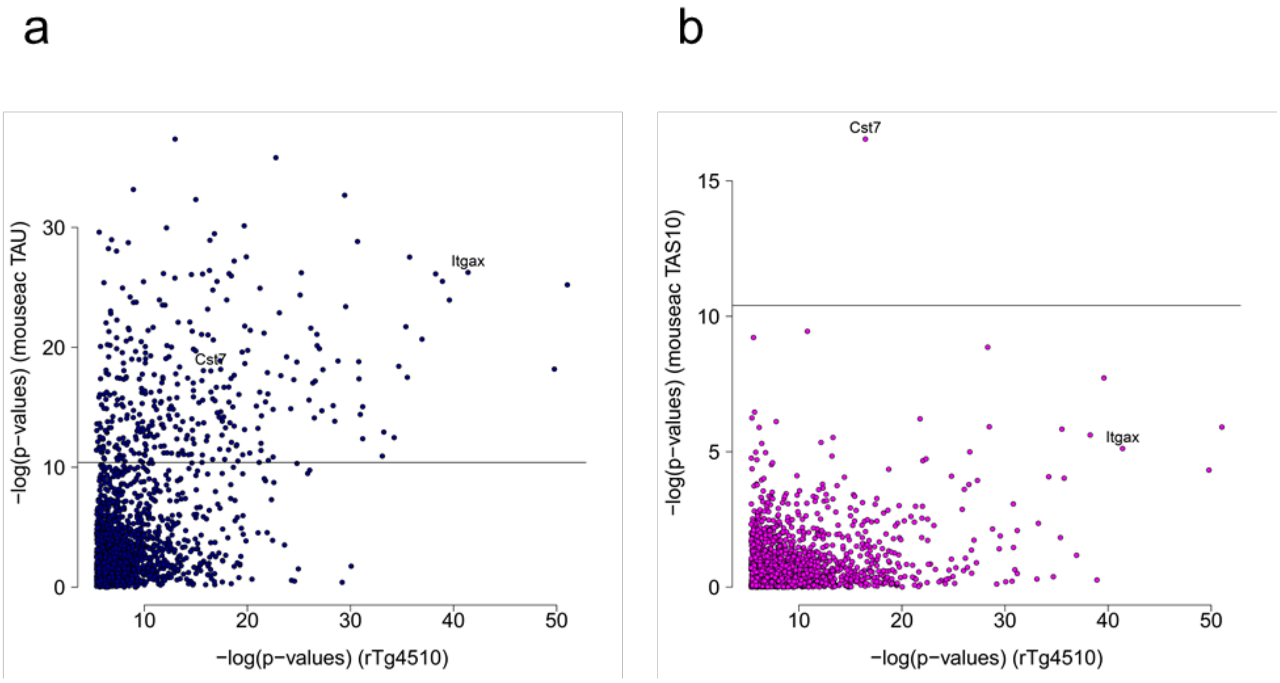

**Supplementary Figure 15. *Cst7* and *Itgax* were similarly altered in both the TAU and TAS10 models.** **(a)** *Cst7* in TAU mice from Mouseac (Total n = 25 animals, 2-4 animals per group, linear regression, $F(3,16) = 48.73$, $\beta = 187.00$, $P = 1.46E-8$). TAU transgenic mice (dark blue) compared to wild type (black) controls. **(b)** *Cst7* in TAS10 mice from Mouseac (Total n = 24 animals, 1-4 animals per group, linear regression, $F(3,15) = 43.42$, $\beta = 11.58$, $P = 6.50E-8$). TAS10 transgenic mice (pink) compared to wild type (black) controls. **(c)** *Itgax* in TAU mice from Mouseac (Total n = 25 animals, 2-4 animals per group, linear regression, $F(3,16) = 137.86$, $\beta = 28.59$, $P = 3.95E-12$). TAU transgenic mice (dark blue) compared to wild type (black) controls. **(d)** *Itgax* in TAS10 mice from Mouseac (Total n = 24 animals, 1-4 animals per group, linear regression, $F(3,15) = 6.03$, $\beta = 0.93$, $P = 0.0060$). TAS10 transgenic mice (pink) compared to wild type (black) controls. Dashed lines represent mean paths across age groups.

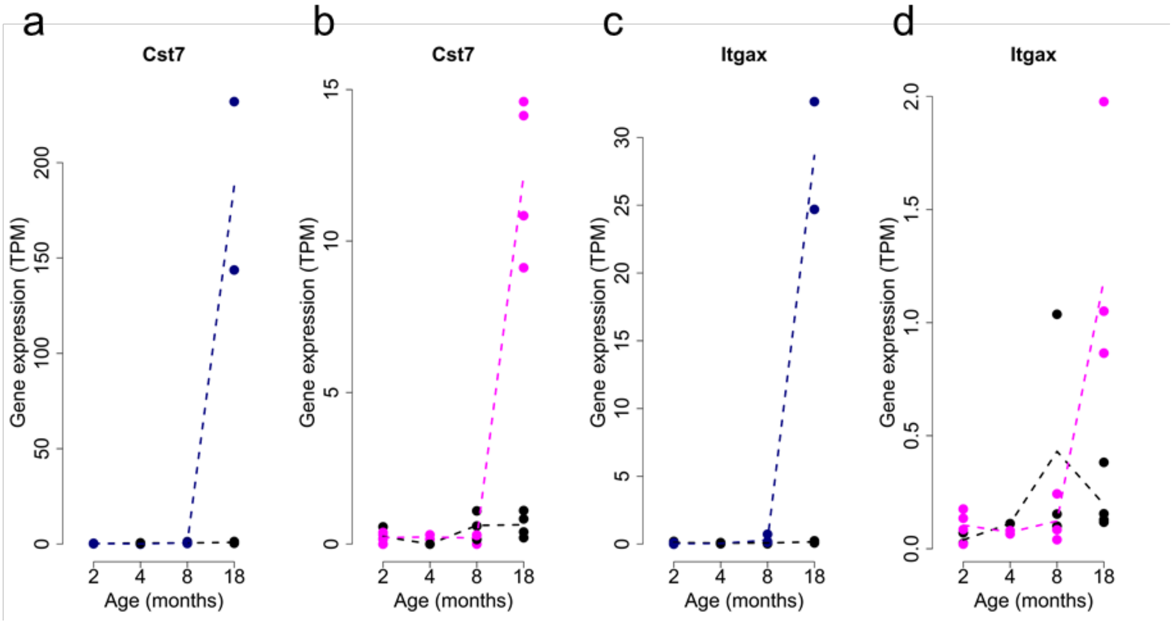

**Supplementary Figure 16. Co-expression modules associated with tau pathology in rTg4510 mice.** In total 18 co-expression modules (each labelled with an arbitrary color) were identified in the rTg4510 TG and WT mice (n = 58 RNA-seq datasets). Shown is the association of each module with genotype, age and progressive changes in TG mice (i.e. the interaction between genotype and age). Bonferroni corrected significant (α=0.05) *P* values are highlighted with an asterisk. The total number of genes in each module is shown in parentheses.

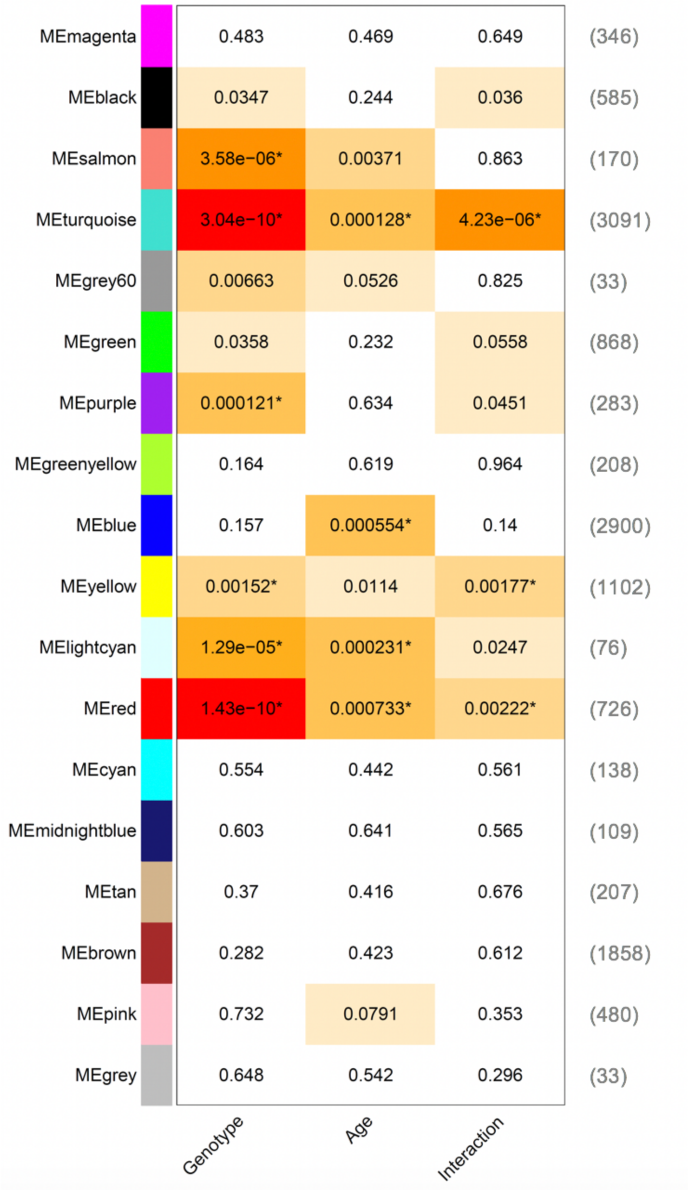

**Supplementary Figure 17. Entorhinal cortex co-expression modules associated with rTg4510 genotype.** Six modules were significantly different (Bonferroni corrected P < 0.05) between TG and WT mice. **(a)** Red module (linear regression, t(53) = -7.93, β = -0.18, P = 1.43E-10). **(b)** Turquoise module (linear regression, t(53) = 7.73, β = 0.18, P = 3.04E-10). **(c)** Salmon module (linear regression, t(53) = 5.17, β = 0.14, P = 3.58E-06). **(d)** Light-cyan module (linear regression, t(53) = -4.81, β = -0.13, P = 1.29E-05). **(e)** Purple module (linear regression, t(53) = -4.15, β = -0.13, P = 0.00012). **(f)** Yellow module (linear regression, t(53) = -3.34, β = -0.10, P = 0.0015). Shown for each module is the module eigengene value for each individual animal (total n = 58 mice). Coloured circles represent rTg4510 TG mice and white circles represent WT control mice. Each circle represents a single mouse. Total n = 58 animals (6-8 animals per group).

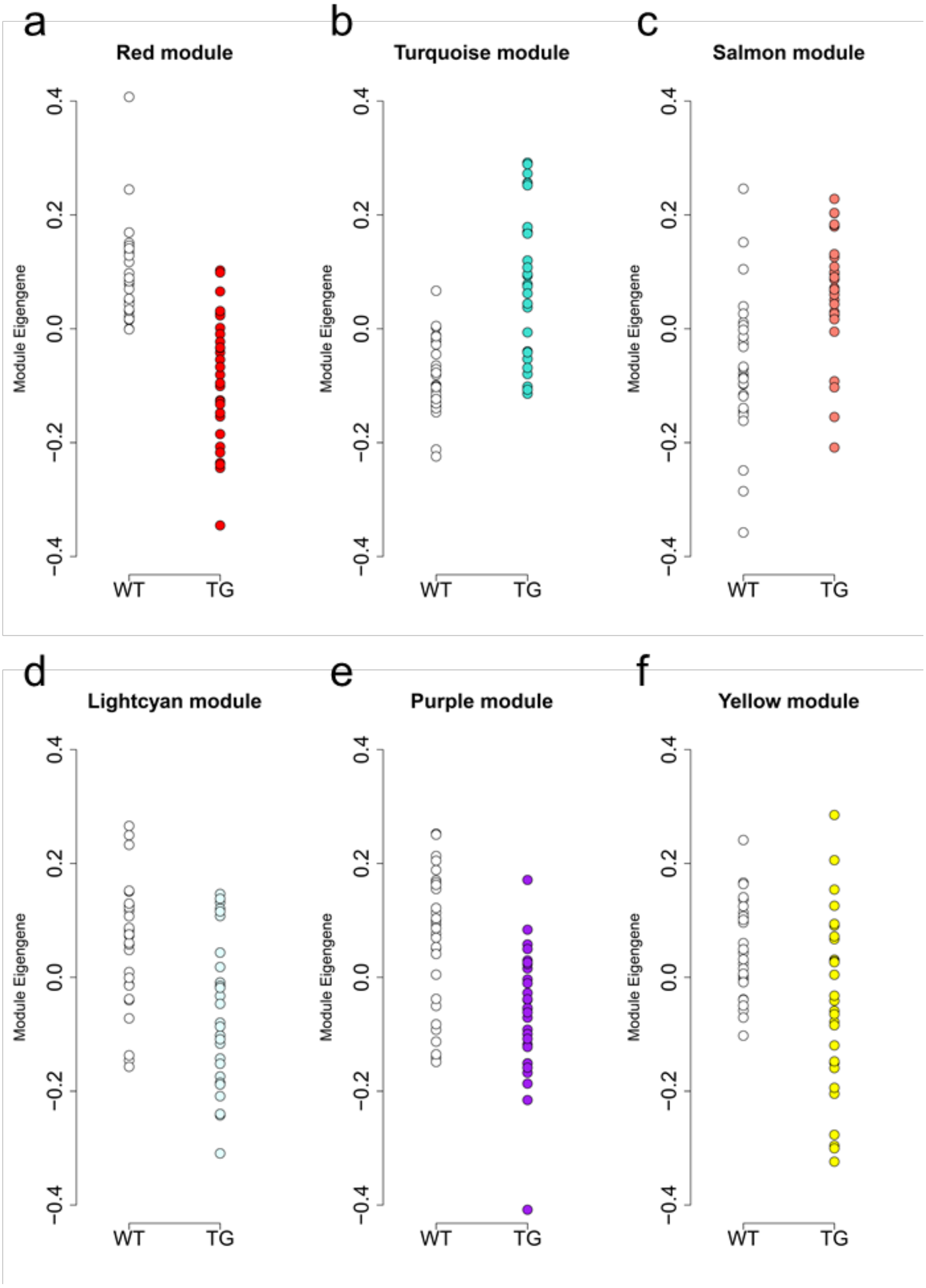

**Supplementary Figure 18. Entorhinal cortex co-expression modules are strongly correlated with tau pathology across multiple brain regions.** Heatmap representing module-trait relationships between module eigengenes and tau pathology measured using immunohistochemistry in each brain region assessed, where red indicates a positive correlation and blue indicates a negative correlation. Values indicate correlation (R) coefficients and their corresponding *P* values in parentheses. Total n = 58 animals.

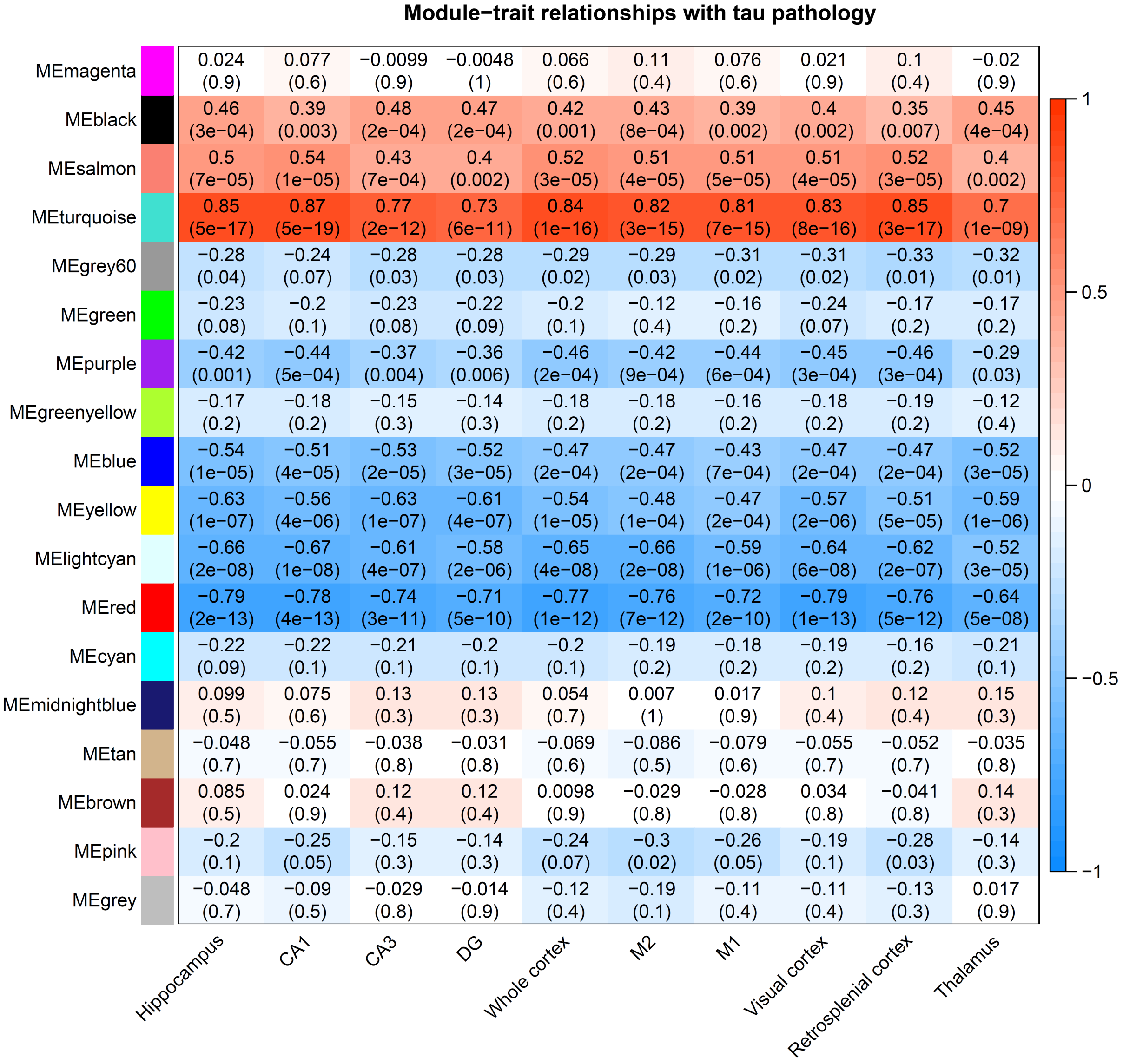

**Supplementary Figure 19. Highly connected transcripts in the three co-expression networks showing progressive changes in rTg4510 mice are most strongly correlated with actual levels of pathology in the hippocampus measured using immunohistochemistry. (a)** Correlation of module membership for each gene with tau pathology in the hippocampus in the turquoise module (n = 3091 transcripts, Pearson correlation, r = 0.85, *P* < 1.00E-200). **(b)** Correlation of module membership for each gene with tau pathology in the hippocampus in the yellow module (n = 1102 transcripts, Pearson correlation, r = 0.49, *P* = 1.30E-67). **(c)** Correlation of module membership for each gene with tau pathology in the hippocampus in the red module (n = 726 transcripts, Pearson correlation, r = 0.80, *P* = 8.88E-163). **(d)** Correlation of intramodular connectivity for each gene with tau pathology in the hippocampus in the turquoise module (Pearson correlation, r = 0.75, *P* < 1.00E-200). **(e)** Correlation of intramodular connectivity for each gene with tau pathology in the hippocampus in the yellow module (Pearson correlation, r = 0.48, *P* = 1.40E-64). **(f)** Correlation of intramodular connectivity for each gene with tau pathology in the hippocampus in the red module (Pearson correlation, r = 0.75, *P* = 4.30E-132). **(g)** Module membership raised to power 6 (y-axis) against intramodular connectivity (x-axis) in turquoise module (Pearson correlation, r = 0.99, *P* < 1.00E-200). **(h)** Module membership raised to power 6 (y-axis) against intramodular connectivity (x-axis) in yellow module (Pearson correlation, r = 0.98, *P* < 1.00E-200). **(i)** Module membership raised to power 6 (y-axis) against intramodular connectivity (x-axis) in red module (Pearson correlation, r = 0.98, *P* < 1.00E-200). Each circle represents a single gene. Total n = 58 animals (6-8 animals per group).

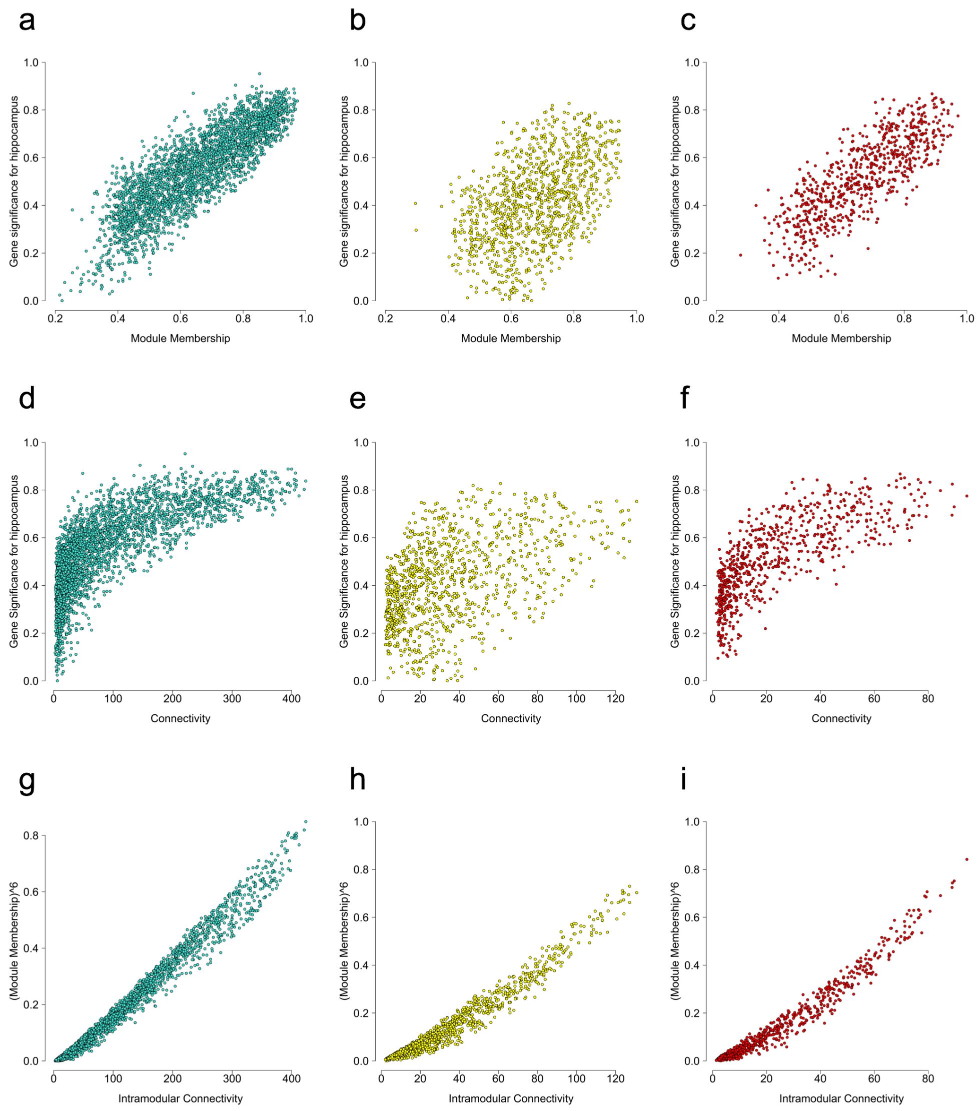

**Supplementary Figure 20. The three tau-associated co-expression modules are most strongly correlated with pathological burden measured by immunohistochemistry in regions of the brain affected early in AD. (a)** Correlation of module eigengenes for each sample with tau pathology in the CA1 subregion of the hippocampus in the turquoise module (Pearson correlation, r = 0.87). **(b)** Correlation of module eigengenes for each sample with tau pathology in the CA3 subregion of the hippocampus in the yellow module (Pearson correlation, r = -0.63). **(c)** Correlation of module eigengenes for each sample with tau pathology in the visual cortex in the red module (Pearson correlation, r = -0.79). **(d)** Correlation of module eigengenes for each sample with tau pathology in the thalamus in the turquoise module (Pearson correlation, r = 0.70). **(e)** Correlation of module eigengenes for each sample with tau pathology in the thalamus in the yellow module (Pearson correlation, r = -0.59). **(f)** Correlation of module eigengenes for each sample with tau pathology in the thalamus in the red module (Pearson correlation, r = -0.64). Correlations and corresponding *P* values for all modules and traits can be seen in **Supplementary Figure 18**. Each circle represents a single individual mouse, with coloured circles representing rTg4510 TG mice and white circles representing WT control mice. Total n = 58 animals (6-8 animals per group).

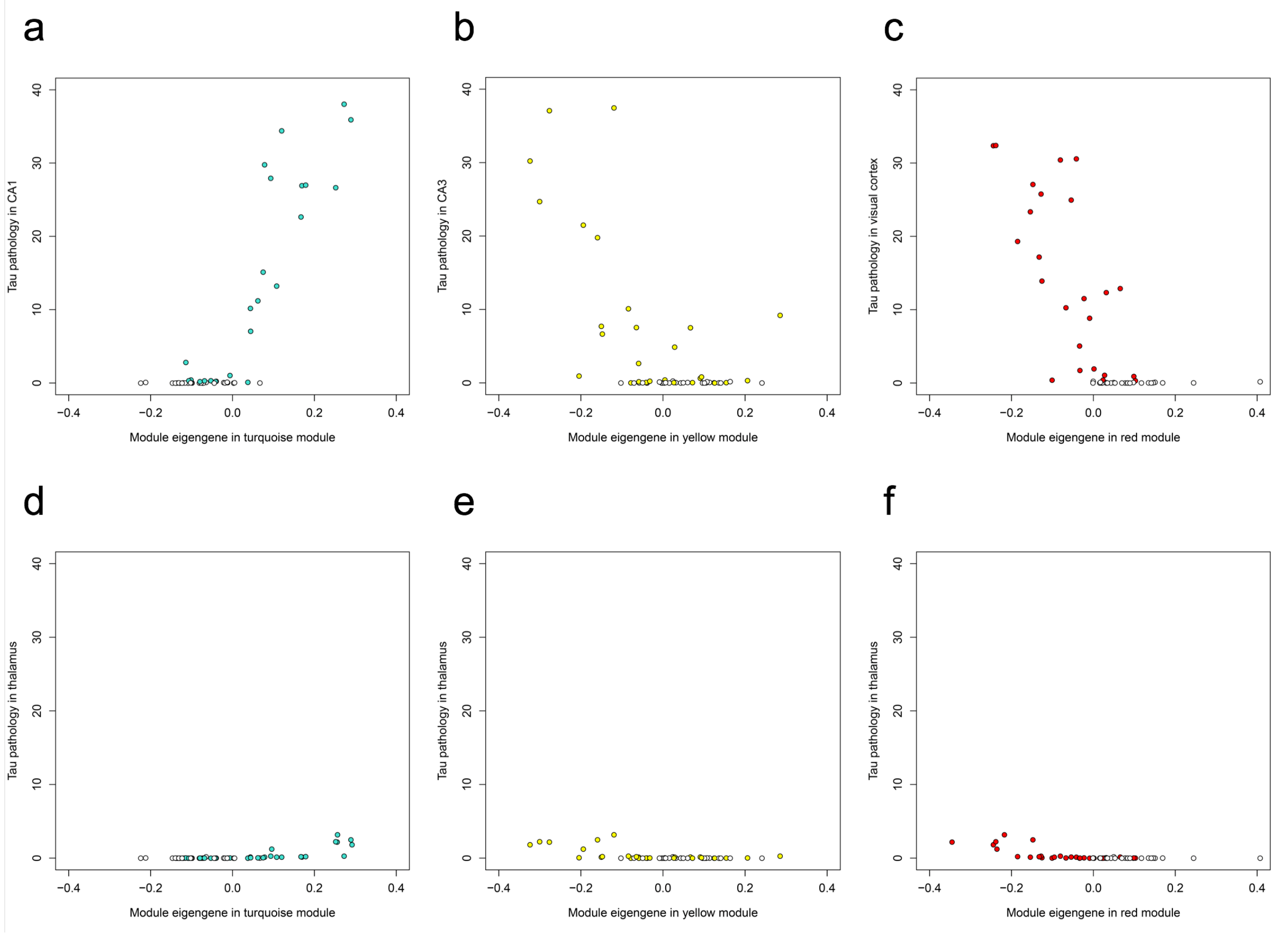

**Supplementary Figure 21. Euclidean distance between samples in rTg4510 mice.** Heatmap of sample-to-sample distances using *rlog*-transformed values.

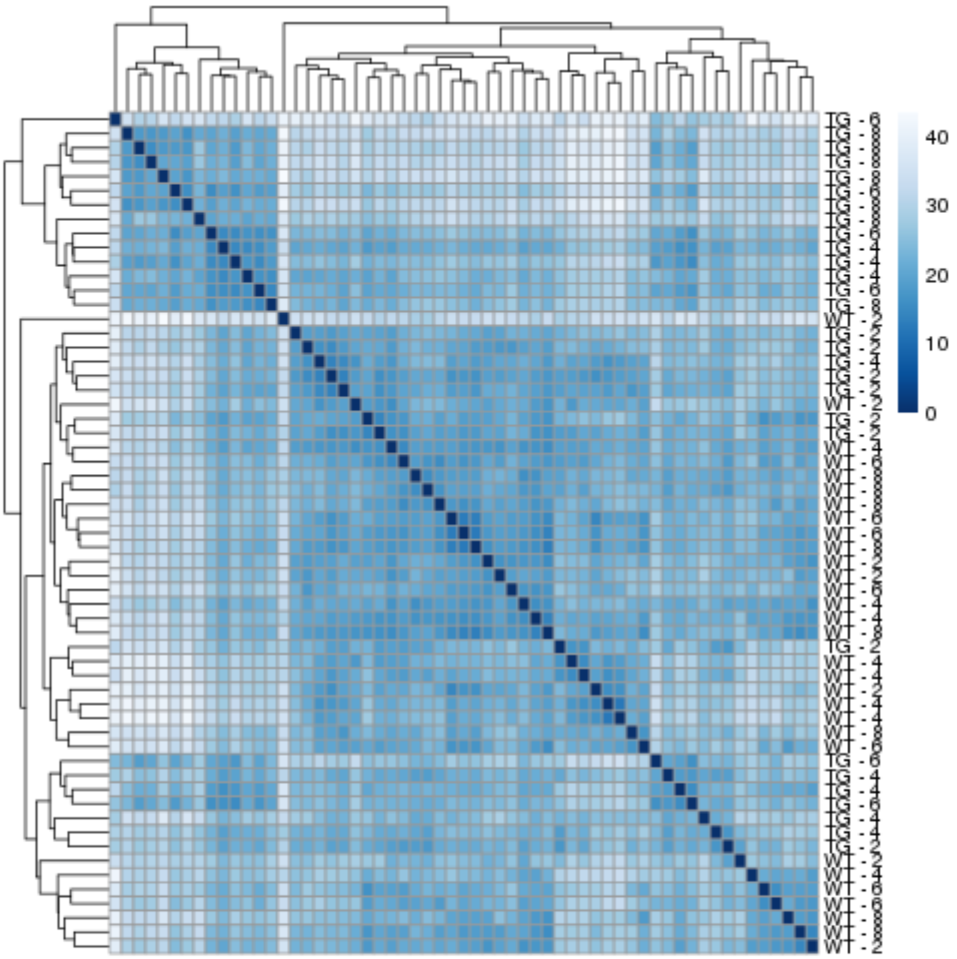

**Supplementary Figure 22. Euclidean distance between samples in J20 mice.** Heatmap of sample-to-sample distances using *rlog*-transformed values.

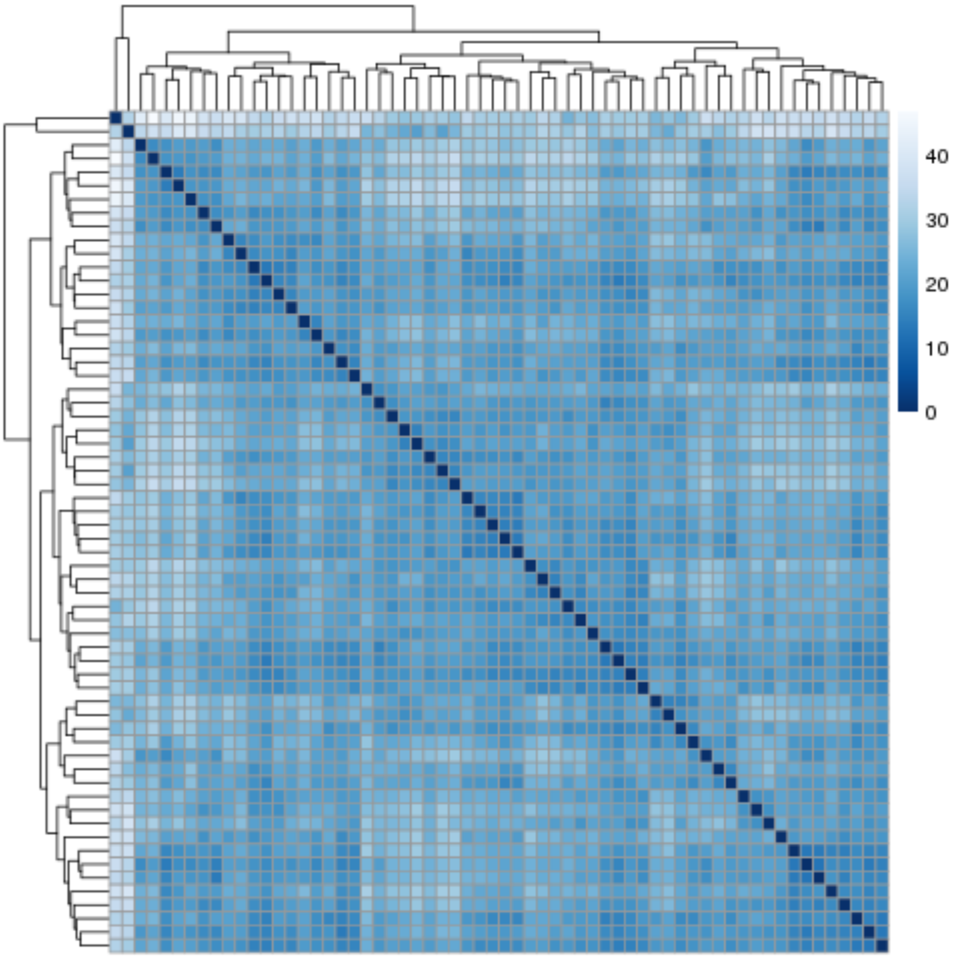

**Supplementary Figure 23. Principal component analysis (PCA) plot of the first two principal components in rTg4510 samples.**

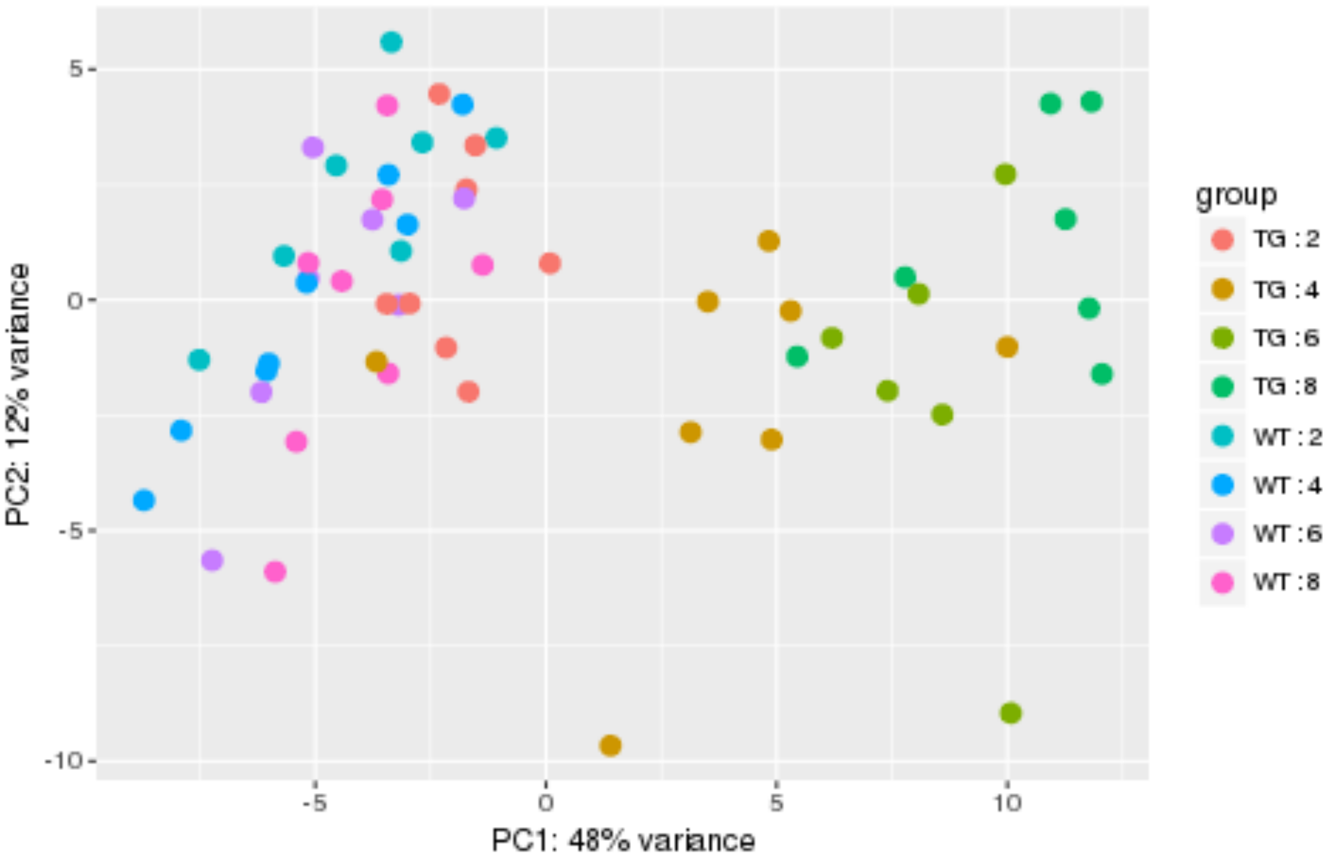

**Supplementary Figure 24. Principal component analysis (PCA) plot of the first two principal components in J20 samples.**

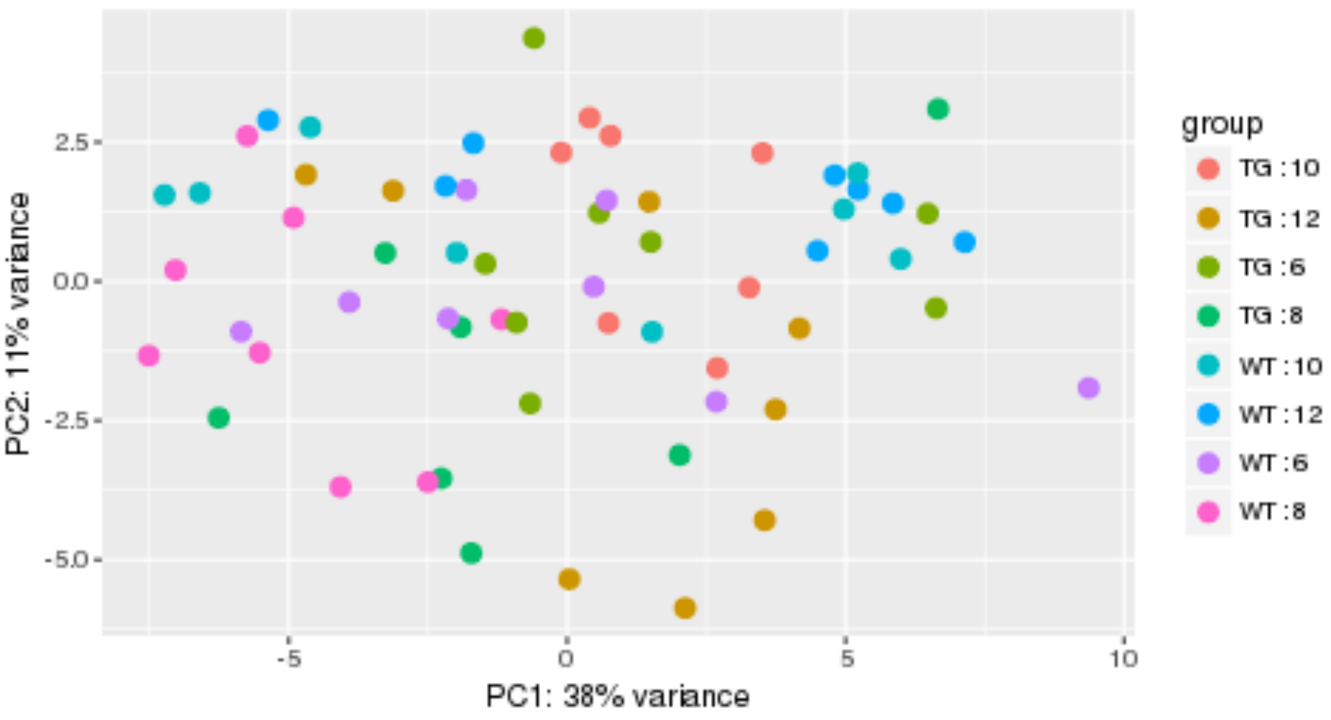
